# Intensive pine plantation management shapes infectious agent communities in the generalist rodent *Apodemus sylvaticus*, with consequences for zoonotic hazard

**DOI:** 10.64898/2026.09.24.753431

**Authors:** Aloïs Berard, Maxime Galan, Julien Pradel, Caroline Tatard, Guillaume Castel, Lily Cesari, Philippe Gauthier, Rachel Beaubaton, Yannick Simonin, Marine Le Guyader, Zouheira Djelouadji, Nathalie Charbonnel

## Abstract

Land use change is widely recognized as a major driver of emerging zoonotic diseases. Yet, while forest plantations are expanding worldwide, their consequences on the circulation of infectious agents in wildlife remain poorly understood.

We investigated how native pine plantations and broadleaved habitats shape rodent-associated infectious hazard in the generalist wood mouse (*Apodemus sylvaticus*), in southwestern France. Using a multi-season survey, we screened 424 wood mice for a broad range of infectious taxa and compared infection probability and infected-host abundance among pine plantation edges, broadleaved hedgerows embedded within pine plantations and broadleaved forests. Using a combination of linear mixed model and joint species distribution modelling, we tested whether infectious hazard differed between pine plantations and broadleaved forests, whether broadleaved hedgerows shifted these patterns towards those observed in broadleaved forests, and how these habitat-related patterns varied across seasons.

We detected 22 potentially infectious taxa, including nine recognized zoonotic bacteria. Season strongly influenced infectious patterns, while habitat effects were taxa and season dependent. In spring, pine plantation edges increased infection probability for *Bartonella grahamii* and *Neoerhlichia mikurensis* compared to broadleaved forest, but their strong negative effect on wood mouse abundance resulted in consistently lower infected-host abundance for most infectious taxa. Broadleaved hedgerows showed distinct and season-dependent patterns with infectious profiles resembling those of pine edges in spring, but with higher infectious hazard than both other habitats for several taxa in autumn.

Our results highlight that habitat effects on infectious hazard cannot be inferred from infection probability alone. Pine plantations may reduce rodent-associated zoonotic hazard through their strong negative effect on host abundance, despite increasing infection probability for some agents. Forest diversification and restoration initiatives, such as broadleaved hedgerows here, may have unforeseen epidemiological consequences, underscoring the need to integrate disease ecology into forest management and restoration to anticipate variation in zoonotic risks, while maximizing ecological benefits.

## 1. Introduction

The burden posed by infectious diseases represents an increasing threat for both biodiversity and human well-being. In wildlife, intensified epizootic events are responsible for substantial mortality and population declines, raising major conservation concerns (Fisher & Garner, 2020; Hoyt et al., 2021). In humans, the frequency of reported infectious disease outbreaks has accelerated over recent decades, with a large proportion originating from wildlife (Jones et al., 2008; Smith et al., 2014; Meadows et al., 2023). This intensification is closely linked to the global changes driven by human activities (Sanderson & Alexander, 2020; Baker et al., 2022; Carlson et al., 2025). Among the multiple and interdependent drivers of disease emergence, historically dominant land-use changes such as deforestation, agricultural expansion, and urbanization are considered major factors (Gottdenker et al., 2014; Fell et al., 2026). Through cascading ecological effects, these changes can reshape infectious hazard and human exposure by reconfiguring host communities, altering host immunity and behaviour, and modifying wildlife-human interactions (Plowright et al., 2021).

Forests are particularly susceptible to environmental changes that can substantially reshape infectious diseases dynamics. They are major terrestrial ecosystems, providing habitat for a large share of terrestrial biodiversity while supporting numerous services (Brockerhoff et al., 2017; Vié et al., 2009). Yet, forests are increasingly affected by human activities, through both widespread deforestation (Singh & Persson, 2026) and alterations of their structure, composition and management, which can substantially erode forest biodiversity (Chaudhary et al., 2016). Such changes may create or exacerbate conditions prone to the outbreak of infectious diseases (Guégan et al., 2020, 2023).

While the extent of natural forests continues to decline, the area covered by forest plantations has increased and is expected to expand further (FAO, 2025). Forest plantations can result from the conversion of natural forests (Ahlström et al., 2022; Singh & Persson, 2026), or be established through the afforestation or reforestation of previously non-forested land (Hua et al., 2018). They typically consist of even-aged stands dominated by one or two tree species (i.e. monocultures), established and managed for wood production (FAO, 2025) or forest recovery purposes (Hua et al., 2018). Forest plantations may influence infectious disease dynamics through multiple, interconnected mechanisms. First, they often support lower wildlife biodiversity than primary and secondary forests, in part by filtering out forest specialists while favouring disturbance-tolerant species (Brockerhoff et al., 2008; Iezzi et al., 2018). Such shifts in animal community composition may have epidemiological consequences as human-dominated ecosystems tend to favour host species that harbour zoonotic pathogens (Gibb et al., 2020). Second, by modifying vegetation structure, microclimate, and the intensity of human disturbances, plantations may further affect host demography (Rubio et al., 2019), vector abundance (Davidson et al., 2024), or pathogen persistence. Lastly, degraded forests and plantations may affect the health status of persisting generalist species, by altering stress levels, diet or gut microbiota, thus potentially modifying their susceptibility to infection (Messina et al., 2018; Heni et al., 2023).

Despite these potential effects, the influence of forest plantations for communities of infectious agents in wildlife remains poorly understood. Evidence from other tree commodity plantations, such as oil palm and rubber, suggests that plantations can increase zoonotic infectious disease risks in nearby human populations (Shah et al., 2019; Morand & Lajaunie, 2021), potentially by altering pathogen transmission or human exposure. Nonetheless, few studies have compared communities of infectious agents between commercial forest plantations and natural forests, and most have focused on a limited number of taxa (Infante et al., 2022; Riquelme et al., 2021), with zoonotic agents rarely receiving attention (but see Rubio et al., 2019). This limited taxonomic scope may hinder our ability to detect general effects of intensive plantation management on disease risks because infectious taxa can respond differently to the same environmental changes, depending on their ecological requirements (Gottdenker et al. 2014; Fell et al. 2026). Moreover, as co-infections are the norm in wildlife (Petney and Andrews 1998), infectious taxa may directly or indirectly interact within hosts (Duncan et al. 2024), potentially affecting transmission dynamics in ways that cannot be captured when studying organisms independently (Telfer et al. 2010; Abbate et al. 2024). Moving beyond these single-pathogen approaches and considering the broader community of infectious taxa circulating within an ecosystem (i.e. the infectiosome; Gasparini 2025) through a multi-taxon perspective is therefore critical to provide a better understanding of how forest plantations shape infectious disease patterns, including zoonotic hazard.

The Landes de Gascogne Forest in southwestern France provides an opportunity to investigate these questions. It is one of the largest intensively managed and human-planted forest landscapes in Europe. Mainly established during the eighteenth and nineteenth centuries on former heathlands and wetlands (Jolivet et al., 2007), it is now dominated by native maritime pine (*Pinus pinaster*), which cover nearly 70% of its total area (Mora et al., 2012). Within this plantation-dominated landscape, linear broadleaved structures persist along pine plantation edges where they are maintained as property boundaries and for their recreational value. Recent studies of these existing broadleaved hedgerows have highlighted local benefits for pest resistance (Plat et al., 2025) and biodiversity (Plat et al., 2026), suggesting that increasing their extent could provide a potential edge-based approach for pine plantation landscape restoration. These structures may therefore offer an opportunity to assess whether increasing broadleaved habitat can counteract some negative ecological consequences of intensive pine plantation management, including potential effects on wildlife-associated infectiosome.

Rodents are particularly relevant in this context. They are the most species-rich mammalian order and consequently host the greatest diversity of zoonotic pathogens among mammals (Han et al., 2015; Mollentze & Streicker, 2020). Their diversity and frequent occurrence in human-modified environments make them useful sentinels for investigating the relationships between land-use changes and epidemiological processes. Among rodents, the generalist wood mouse *Apodemus sylvaticus* is particularly well suited to this question because of its abundance and widespread distribution across European forests as well as its ability to persist in various habitats, from relatively undisturbed forests and forest plantations, to agricultural, peri-domestic and urban environments (Tattersall et al., 2001; Pradel et al., 2022; Berard et al., 2026). In addition, *A. sylvaticus* is involved in the maintenance and the circulation of multiple zoonotic pathogens (Fabbri, Dorigo, et al., 2026; Fabbri, Grassi, et al., 2026). Previous work in the Landes de Gascogne landscape further showed that wood mouse abundance varies across habitat types and season (Berard et al., 2026), with broadleaved structures supporting higher abundances than pine plantations, and autumn characterized by higher abundances than spring. Whether such habitat-related differences in host abundance are accompanied by changes in infectious agent community composition, richness, and prevalence remains unknown.

In this study, we assessed how pine-plantation-dominated landscapes and broadleaved hedgerows shape the infectiosome of *Apodemus sylvaticus*, the dominant rodent species in the Landes de Gascogne Forest (Berard et al., 2026), using a multi-year, multi-season and multi-taxon survey. Specifically, we aimed to (i) characterize the wood mouse infectiosome across this forested landscape and identify potential zoonotic taxa; (ii) test whether infectiosome patterns in intensively managed pine plantations differ from those observed in broadleaved forests; and (iii) assess whether broadleaved hedgerows modify these patterns within the pine-dominated landscape. Given the intensive management of pine plantations, we hypothesized that these habitats would provide conditions favouring infectious agents transmission, thus increasing infection probabilities overall. At the same time, their negative effect on host abundance was predicted to result in contrasting infected-host abundance trends across infectious taxa, depending on the extent to which the infection probability offset reduced host abundance. Conversely, because hedgerows provide unmanaged broadleaved vegetation within the pine matrix, we expected them to attenuate these differences, leading to infectiosome characteristics more similar to those observed in broadleaved forests. We further predicted these patterns to vary across seasons, given the marked seasonal variation in host abundance.

## 2. Methods

### 2.1 Ethical statement

The CBGP laboratory is authorized by the Departmental Directorate for Population Protection (DDPP, Hérault, France; approval no. F-34-169-001) for the capture of small mammals and the storage and use of their tissues. All procedures complied with French and European regulations governing the use of animals for scientific purposes (Directive 2010/63/EU). All procedures have undergone validation by the regional ethics committee Comité d’Ethique pour l’Expérimentation Animale Languedoc Roussillon n°36 (approval no. 2073, 2023).

### 2.2 Apodemus sylvaticus sampling

Rodents were sampled following a standardized trapping protocol consisting of three to four consecutive trapping nights with 20 traps per line during five field sessions between 2023 and 2025, including three late spring (May-June) and two autumn sessions (September-October). The study area encompassed 53 sampling sites distributed across 1,050 km² in southwestern France (Figure 1; centre: 44°33’38.2"N, 0°46’36.7"W), with sites classified into three habitat types: edges of monospecific pine plantation (12 sites), broadleaved hedgerows embedded within the pine plantation landscape (24 sites), and broadleaved forests (17 sites), as described in Berard et al. (2026). Broadleaved forest sites were added to the sampling design from 2024 onwards.

**Figure 1.**
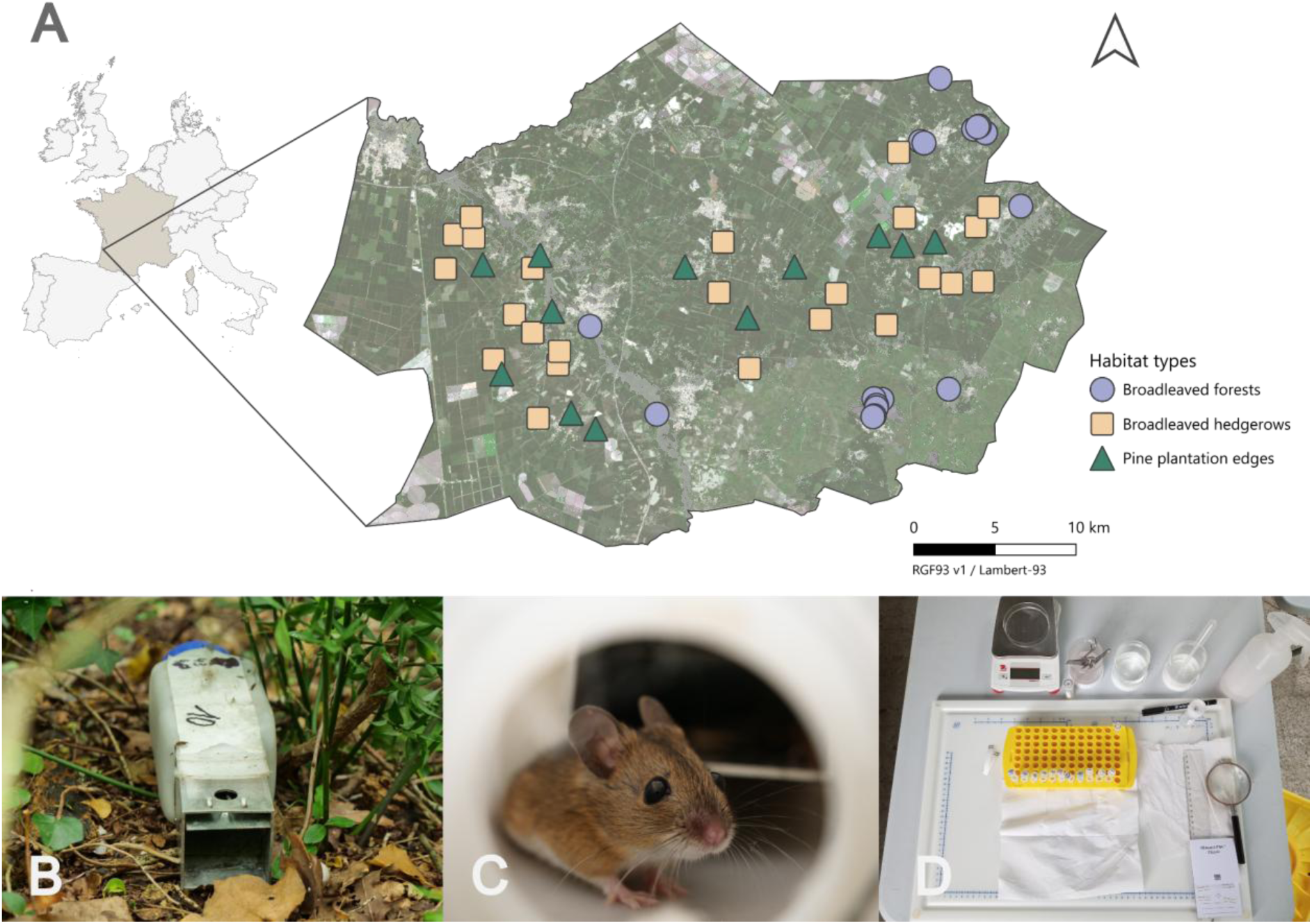
Study area and field sampling procedures. **A.** Distribution of the sampling sites within the Landes Forest (southwestern France). The background combines satellite imagery and coniferous forest cover layer produced by CNES for the Theia Data Centre using Copernicus products. **B.** INRA live trap used for rodent capture. **C.** Live-trapped *Apodemus sylvaticus* individual. **D.** Tissue collection for infectious agent screening.

Individuals were anaesthetized using isoflurane and euthanized by cervical dislocation as recommended by Mills et al. (1995). They were weighed, sexed and measured. Dried blood spot, kidney and spleen samples were collected immediately thereafter. Kidney and spleen samples were preserved in ethanol for subsequent analyses. Rodent species identification relied on morphological characteristics, with molecular confirmation performed using COI barcoding (Galan et al., 2018) or arbitrarily primed polymerase chain reaction (AP-PCR; Bugarski-Stanojević et al., 2008) when morphological assignment was uncertain. Individuals were classified as sexually mature or immature according to external and internal reproductive characteristics (Supplementary Figure S1).

In accordance with ethical guidelines, rodents captured beyond the authorized removal number were marked and released in order to avoid recapture bias. Local *A. sylvaticus* abundance was indexed by the number of unique individuals captured during the first three trapping nights at each site and sampling session, following Spitz et al. (1974).

### 2.3 Infectious agent screening

Infectious status of *A. sylvaticus* individuals was investigated using targeted serological and molecular assays, combined with metabarcoding approaches used to characterize blood-circulating bacteria communities without prior assumptions.

#### 2.3.1 Serological assays for viral exposure

Resuspended dried blood spot samples were analysed by sandwich enzyme linked immunosorbent assays (ELISAs) to detect immunoglobulin G (IgG) antibodies as indicators of previous viral exposure. Antibodies specific to, or cross-reacting with Tula (TULV) and Hantaan (HTNV) orthohantaviruses were assessed using *Orthohantavirus tulaense* and recombinant *Orthohantavirus hantanense* antigens (HTV-001, Prospec), while orthoflavivirus exposure was evaluated using a commercial pan-orthoflavivirus ELISA (Dubois et al., 2017; Beaubaton et al., 2025).

#### 2.3.2 Molecular detection of pathogenic Leptospira spp

Infection with pathogenic *Leptospira* spp. was investigated using DNA extracted from kidney samples (BioBasic EZ-10 Genomic DNA kit, Animal Sample) and a quantitative real-time PCR (qPCR) assay targeting the *lipL32* gene (Dobigny et al., 2015 adapted from Stoddard et al., 2009). All qPCR reactions were performed in duplicate on a LightCycler 480 (Roche). Samples yielding a cycle threshold (Ct) value < 40 for both duplicates were considered positive. Each run included a positive and a negative control. Subsequently, we characterized *Leptospira* species and serovars. All *lipL32-*positive samples were typed using the *lfb1* gene following the protocol described in Garcia-Lopez et al. (2024). For further statistical analyses, only samples positive for both *lipL32* and *lfb1* assays were considered infected with pathogenic *Leptospira* spp. and the occurrences of the *Leptospira* spp. taxa were pooled into a single *Leptospira* spp. taxon.

#### 2.3.3 Molecular detection of bacteria using 16S rRNA metabarcoding

We applied the metabarcoding approach described in Galan et al. (2016) to detect additional potentially infectious bacteria from spleen samples. This organ was used because its blood-filtering and reserve functions make it suitable for detecting circulating microorganisms. DNA was extracted using the DNeasy® 96 Blood & Tissue kit (Qiagen). The V4 hypervariable region of the 16S rRNA gene was amplified and sequenced using Illumina MiSeq (2 x 251 pb). All DNA extractions were amplified by PCR and sequenced twice independently using two different libraries for each sample. Negative controls for DNA extraction, PCR amplification, and indexing, as well as mock bacterial communities were included throughout the complete workflow.

Sequences were analysed using the FROGS pipeline (Find Rapidly OTU with Galaxy Solution; Escudié et al., 2018). Reads were clustered into high-resolution operational taxonomic units (OTUs) using SWARM (fastidious method with *d = 1*) and taxonomically assigned based on reference sequences from the SILVA database (138.2 Pintail100) using BLASTN+. SILVA annotations were subsequently updated according to the current LPSN recommendations (Göker et al. 2025). Additional filtering steps following (Galan et al., 2016) were applied to remove potential false-positives detections. Unassigned OTUs were manually inspected using the NCBI nucleotide database, allowing the identification of apicoplast-derived 16S rRNA sequences belonging to Apicomplexan parasites (Zintl et al., 2023). Mycoplasmataceae-affiliated OTUs were further investigated using BLAST searches and phylogenetic analyses, with taxonomic identification following the classification of Gupta et al. (2018).

For OTUs assigned to bacterial taxa, only the ones identified at least to the genus level and representing more than 0.001% of total reads were retained. OTUs assigned to the same taxon were merged at the highest reliable taxonomic resolution. Taxa typically associated with environmental or gut microbiota were excluded.

#### 2.3.4 Species characterization of Bartonella spp. using gltA and rpoB metabarcoding

To refine identification of *Bartonella* spp. detected by 16S rRNA metabarcoding, all *Bartonella*-positive samples were further analysed using a targeted metabarcoding approach based on the *gltA* and *rpoB* genes (primers from Norman et al., 1995; Renesto et al., 2001; Supplementary Methods). Sequencing, filtering and clustering were performed as described above. Taxonomic assignment of *rpoB* OTUs was conducted using the rpoB_INRA_122017 database (Ogier et al., 2019). Taxonomic assignment of all *gltA* OTUs, as well as *rpoB* OTUs showing less than 97% sequence identity or lacking taxonomic assignment, was investigated using the NCBI nucleotide database. Only OTUs identified at the species level and representing more than 0.01% of total reads were retained. For each gene, OTUs assigned to the same species were merged. Species detections obtained from the two markers were then combined, regardless of whether a species was detected by one or both markers.

### 2.4 Statistical analyses

For subsequent analyses, infectious taxa were considered as present or absent in each individual, irrespective of the detection method or sequence read abundance.

Three complementary modelling approaches were used to investigate variations in the infectiosome. First, we analysed the factors driving individual-level infectious taxon richness. Then, we examined pathogen-specific responses using two Joint Species Distribution Models (JSDMs), focusing respectively on i) the probability of infection at the individual wood mouse level and ii) the abundance of infected wood mice at the site-session level.

Data exploration and analyses were performed with R Statistical Software (v4.6.0), except for JSDMs, which were run on the genotoul bioinformatics platform Toulouse Occitanie (R module v4.5.0).

#### 2.4.1. Individual-level infectious taxon richness

We first investigated the drivers of variation in infectious taxon richness at the individual host level using generalized linear mixed models (GLMMs) with a Poisson error distribution. Richness was defined as the number of distinct taxa detected per individual. We evaluated the effects of three individual host characteristics (weight, sex and sexual maturity) and three site-session characteristics (habitat type: pine edge, broadleaved hedgerow or broadleaved forest; season: spring or autumn; and local *A. sylvaticus* abundance). Because individual characteristics were missing for two wood mice, these individuals were excluded from individual-level analyses.

We built a set of six models, representing alternative biological hypotheses on the determinants of richness (Supplementary Table S1). The null model assumed no effect of any predictor. The host model tested whether individual characteristics alone explained variation in richness. Two additional models evaluated the independent contribution of habitat type and season, respectively, while controlling for host characteristics and local wood mouse abundance. Finally, two models tested whether habitat and season jointly explained additional variation through additive effects or through their interaction. Sampling site identity, individual identity and year were included as random intercepts in all models.

Models were compared using an information theoretic approach based on the corrected Akaike information criterion (AICc). The model with the lowest AICc was retained for inference. To interpret the effects of the interaction between habitat and season, post-fit comparisons were conducted on response-scaled estimated marginal means using Tukey’s range test (α = 0.05).

Models were fitted using the package glmmTMB (v1.1.14), with residual diagnostics performed using DHARMa (v0.5.0) and marginal means estimated using emmeans (v2.0.3).

#### 2.4.2 Infectious taxon-specific responses

To evaluate taxon-specific responses of infectious agents to habitat type and season, we applied JSDMs, which simultaneously model the responses of multiple taxa to environmental covariates while accounting for residual dependence among taxa (Warton et al., 2015). Specifically, we used the Hierarchical Modelling of Species Communities (HMSC) framework, a Bayesian generalised linear latent variable model that enables taxon-specific responses to environmental covariates to be related to species traits and phylogenetic relationships (Ovaskainen et al. 2017).

The first HSMC approach was designed to examine infection probability at the individual level by modelling the presence/absence of each infectious taxon in individual hosts using a probit distribution. The second HSMC approach modelled the abundance of infected individuals at the site-session level using a lognormal Poisson distribution to account for overdispersion. For both approaches, we compared the same set of hypothesis-driven models as those previously described for infectious taxon richness (Supplementary Table S1).

To ensure sufficient statistical power, infectious taxa detected in fewer than 15 individuals were excluded, leaving 12 taxa. For the abundance analysis, only individuals caught during the first three trapping nights were kept in order to standardize trapping effort among site-sessions, thus excluding three individuals. Because not all captured rodents were screened for infectious agents, the five site-sessions containing more than one unscreened individual were also excluded, thereby limiting potential bias due to unequal screening coverage among site-sessions.

For all models, field session and site identity were included as unstructured random effects, with site identity also included as a spatially structured random effect. Individual identity was included as an additional random effect in the infection probability models. We incorporated phylogenetic relationships among infectious taxa, inferred from their taxonomic classification, to test whether closely related taxa exhibited more similar responses to fixed covariates than distantly related ones. We also included two binary transmission traits indicating whether each taxon was known to be vector-borne and/or transmissible through non-vector pathways (i.e. direct or environmental transmission).

HMSC models were fitted using the Hmsc package (v3.3-7; Tikhonov et al., 2020) with default prior distributions. Posterior distributions were sampled using four Markov Chain Monte Carlo (MCMC) chains, each run for 1,200,000 iterations, with the first 400,000 iterations discarded as burn-in. A thinning interval of 1,000 iterations was applied, yielding 800 posterior samples per chain. Model convergence was assessed using the potential scale reduction factor (PSRF; Gelman & Rubin, 1992).

Model performance was assessed in terms of explanatory and predictive power. For infection probability models, performance was evaluated using the area under the receiver operating characteristic curve (AUC) and Tjur’s coefficient of discrimination (Tjur’s R²). For abundance models, performance was assessed using the pseudo-*R*² based on the squared Spearman correlation (SR²). Performance metrics corresponded to the mean values across taxa. Explanatory power was estimated from predictions based on models fitted to the full dataset, whereas predictive power was computed by performing five-fold cross-validation. Models with the highest explanatory Tjur’s R² and SR² values were selected.

This HMSC approach provided estimates of taxon-specific regression coefficients describing responses to fixed covariates (β), as well as parameters quantifying trait effects (Γ), phylogenetic signal (ρ) and residual associations among taxa (Ω). The strength of support for all these parameters was assessed using posterior support (PS), defined as the proportion of posterior samples having the same sign (positive or negative) as the posterior mean. Parameters with PS ≥ 0.95 were considered strongly supported. We also used variance partitioning to assess the relative contributions of fixed and random effects to the variance of response variables for each taxon.

To interpret habitat by season interactions, we generated marginal response predictions for each habitat-season combination. Predictions were obtained for each posterior MCMC sample, allowing pairwise comparisons among habitat x season combinations to be computed from the posterior predictions. Effect sizes and posterior support were then estimated from these contrasts. We applied the same prediction and comparison framework, to investigate trait effects using posterior predictions of community-weighted mean (CWM) as the response variable.

## 3. Results

### 3.1 Composition and diversity of the wood mouse infectiosome

A total of 474 *Apodemus sylvaticus* individuals were captured across the 53 sampling sites, of which 424 were screened for all infectious agents included in the study (Supplementary Table S2).

None of the individuals tested by serological assay were positive for antibodies against targeted *Orthohantavirus* or *Orthoflavivirus*, indicating no evidence of previous exposure to these viruses.

Molecular screening for pathogenic *Leptospira* spp. detected 19 positive individuals (4.5%). Further analyses identified *Leptospira interrogans* (genovar Australis) as the infecting species in 17 individuals and *L. borgpetersenii* (genovar Ballum, Javanica, or Mini) in one individual, whereas one positive sample remained unresolved at the species level.

The 16S rRNA metabarcoding approach enabled the detection of bacterial taxa belonging to several genera of potential infectious relevance, including *Bartonella* spp. (56.3%; n = 239), *Eperythrozoon* spp. (syn. *Mycoplasma*, *Haemobartonella*; 37.9%; n = 161), *Neoehrlichia mikurensis* (15.3%; n = 65) and other less prevalent taxa (*Borrelia, Ehrlichia, Streptobacillus)* together with various Mycoplasmatota taxa. Phylogenetic analyses of Mycoplasmataceae-affiliated sequences resolved 15 OTUs into eight species-level taxa and three phylogenetic clusters that were retained for subsequent analyses (Supplementary Figure S2). In addition, two OTUs corresponding to apicoplast DNA were detected, revealing the presence of apicomplexan parasites from the family Sarcocystidae.

Targeted *gltA* and *rpoB* metabarcoding further resolved *Bartonella* sequences into five species: *B. taylorii*, *B. birtlesii*, *B. grahamii*, *B. doshiae*, and *B. elizabethae*. *Bartonella* remained among the most prevalent infectious-associated bacterial taxa, even after genus-level assignments were refined to species level.

Altogether, the infectiosome characterized here in *A. sylvaticus* comprised 22 potentially infectious taxa. Among the 15 taxa resolved to species level, nine corresponded to recognized zoonotic agents (Table 1).

**Table 1.**
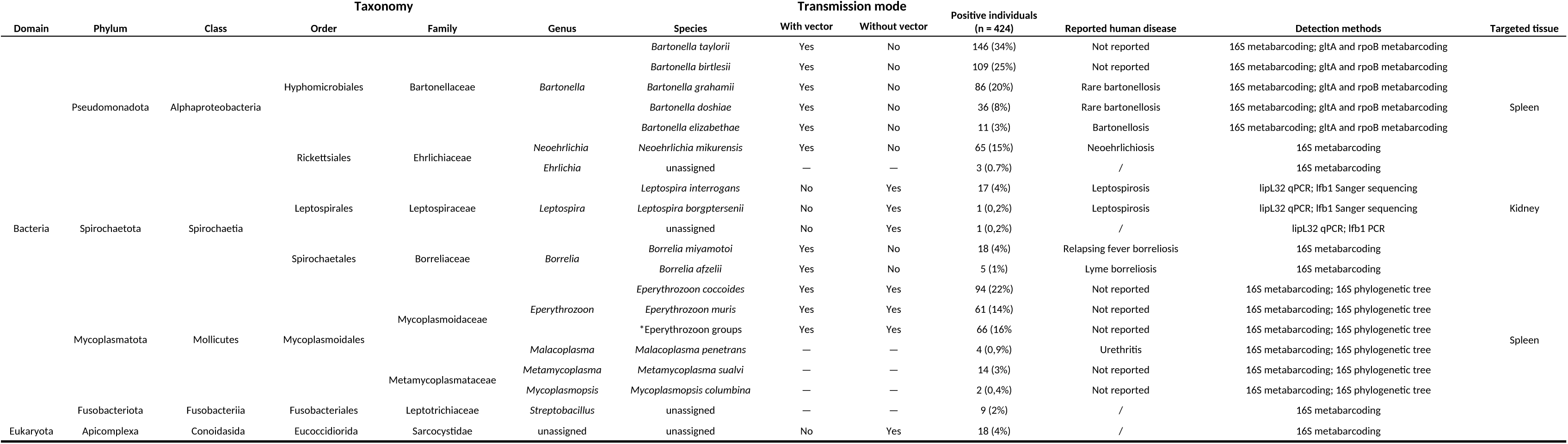
Infectious taxa detected in *Apodemus sylvaticus*. Bacterial nomenclature follows the LPSN (Göker et al., 2025), except for members of the Mycoplasmatota phylum, for which the classification proposed by Gupta et al. (2018) was used. Taxa are reported according to the best taxonomic resolution achieved with lower unresolved ranks indicated as "unassigned". According to the current LPSN nomenclature, *Eperythrozoon coccoides* and *Eperythrozoon muris* are respectively currently recognized as *Mycoplasma coccoides* and *Haemobartonella muris*. *Eperythrozoon OTUs that could not be confidently assigned to a species were grouped into three clusters. Association with human diseases is specified only for taxa identified to the species level, with taxa not identified to species level indicated by “/”. The notation “Not reported” indicates that no cases have been reported for this species to date. Transmission modes that could not be inferred from the species identity or taxonomy are indicated by “—“.

| Taxonomy |  |  |  |  |  |  | Transmission mode |  |  |  |  |  |
|---|---|---|---|---|---|---|---|---|---|---|---|---|
| Domain | Phylum | Class | Order | Family | Genus | Species | With vector | Without vector | Positive individuals (n = 424) | Reported human disease | Detection methods | Targeted tissue |
| Bacteria | Pseudomonadota | Alphaproteobacteria | Hyphomicrobiales | Bartonellaceae | *Bartonella* | *Bartonella taylorii* | Yes | No | 146 (34%) | Not reported | 16S metabarcoding; gltA and rpoB metabarcoding | Spleen |
|  |  |  |  |  |  | *Bartonella birtlesii* | Yes | No | 109 (25%) | Not reported | 16S metabarcoding; gltA and rpoB metabarcoding |  |
|  |  |  |  |  |  | *Bartonella grahamii* | Yes | No | 86 (20%) | Rare bartonellosis | 16S metabarcoding; gltA and rpoB metabarcoding |  |
|  |  |  |  |  |  | *Bartonella doshiae* | Yes | No | 36 (8%) | Rare bartonellosis | 16S metabarcoding; gltA and rpoB metabarcoding |  |
|  |  |  |  |  |  | *Bartonella elizabethae* | Yes | No | 11 (3%) | Bartonellosis | 16S metabarcoding; gltA and rpoB metabarcoding |  |
|  |  |  | Rickettsiales | Ehrlichiaceae | *Neoehrlichia* | *Neoehrlichia mikurensis* | Yes | No | 65 (15%) | Neoehrlichiosis | 16S metabarcoding |  |
|  |  |  |  |  | *Ehrlichia* | unassigned | — | — | 3 (0.7%) | / | 16S metabarcoding |  |
|  | Spirochaetota | Spirochaetia | Leptospirales | Leptospiraceae | *Leptospira* | *Leptospira interrogans* | No | Yes | 17 (4%) | Leptospirosis | lipL32 qPCR; lfb1 Sanger sequencing | Kidney |
|  |  |  |  |  |  | *Leptospira borgptersenii* | No | Yes | 1 (0,2%) | Leptospirosis | lipL32 qPCR; lfb1 Sanger sequencing |  |
|  |  |  |  |  |  | unassigned | No | Yes | 1 (0,2%) | / | lipL32 qPCR; lfb1 PCR |  |
|  |  |  | Spirochaetales | Borreliaceae | *Borrelia* | *Borrelia miyamotoi* | Yes | No | 18 (4%) | Relapsing fever borreliosis | 16S metabarcoding | Spleen |
|  |  |  |  |  |  | *Borrelia afzelii* | Yes | No | 5 (1%) | Lyme borreliosis | 16S metabarcoding |  |
|  | Mycoplasmatota | Mollicutes | Mycoplasmoidales | Mycoplasmoidaceae | *Eperythrozoon* | *Eperythrozoon coccoides* | Yes | Yes | 94 (22%) | Not reported | 16S metabarcoding; 16S phylogenetic tree |  |
|  |  |  |  |  |  | *Eperythrozoon muris* | Yes | Yes | 61 (14%) | Not reported | 16S metabarcoding; 16S phylogenetic tree |  |
|  |  |  |  |  |  | *Eperythrozoon groups | Yes | Yes | 66 (16% | Not reported | 16S metabarcoding; 16S phylogenetic tree |  |
|  |  |  |  |  | *Malacoplasma* | *Malacoplasma penetrans* | — | — | 4 (0,9%) | Urethritis | 16S metabarcoding; 16S phylogenetic tree |  |
|  |  |  |  | Metamycoplasmataceae | *Metamycoplasma* | *Metamycoplasma sualvi* | — | — | 14 (3%) | Not reported | 16S metabarcoding; 16S phylogenetic tree |  |
|  |  |  |  |  | *Mycoplasmopsis* | *Mycoplasmopsis columbina* | — | — | 2 (0,4%) | Not reported | 16S metabarcoding; 16S phylogenetic tree |  |
|  | Fusobacteriota | Fusobacteriia | Fusobacteriales | Leptotrichiaceae | *Streptobacillus* | unassigned | — | — | 9 (2%) | / | 16S metabarcoding |  |
| Eukaryota | Apicomplexa | Conoidasida | Eucoccidiorida | Sarcocystidae | unassigned | unassigned | No | Yes | 18 (4%) | / | 16S metabarcoding |  |

### 3.2 Drivers of individual-level infectious taxon richness revealed by GLMM

The majority of individuals were positive for at least one potentially infectious taxon (75%; n = 318), with nearly half carrying multiple taxa (49%; n = 207). Co-infected individuals harboured an average of 3.2 taxa. In particular, co-infections with multiple *Bartonella* species within individual hosts were common, with 46.8% (n = 112) of *Bartonella*-positive individuals carrying more than one species and an average of 2.3 *Bartonella* species per co-infected host.

The best-supported GLMM explaining infectious taxon richness included host characteristics, season, habitat type, and the interaction between season and habitat (GLMM_6_; AICc = 1333; df = 13; Supplementary Table S3).

Individual infectious taxon richness increased with body mass (β = 0.092; 95% CI = [0.072, 0.112]; *p* < 0.001) and was higher in males than females (β = 0.157; CI = [0.007, 0.308]; *p* = 0.040).

Across habitats, infectious taxon richness differed between seasons, with higher values in late spring than in autumn (estimated marginal mean ratio, R = 0.53, 95% CI = [0.408, 0.688], Tukey-adjusted *p* < 0.001; Figure 2). Pairwise comparisons among habitats within each season revealed no significant differences after Tukey correction (Figure 2). Nevertheless, in autumn, infectious taxa richness tended to be higher in hedgerows than in broadleaved forests (R = 1.50, 95% CI = [0.97, 2.32], *p* = 0.080), whereas in spring it tended to be higher at pine edges than in broadleaved forests (R = 1.32, 95% CI = [0.54, 1.08], *p* = 0.154).

**Figure 2.**
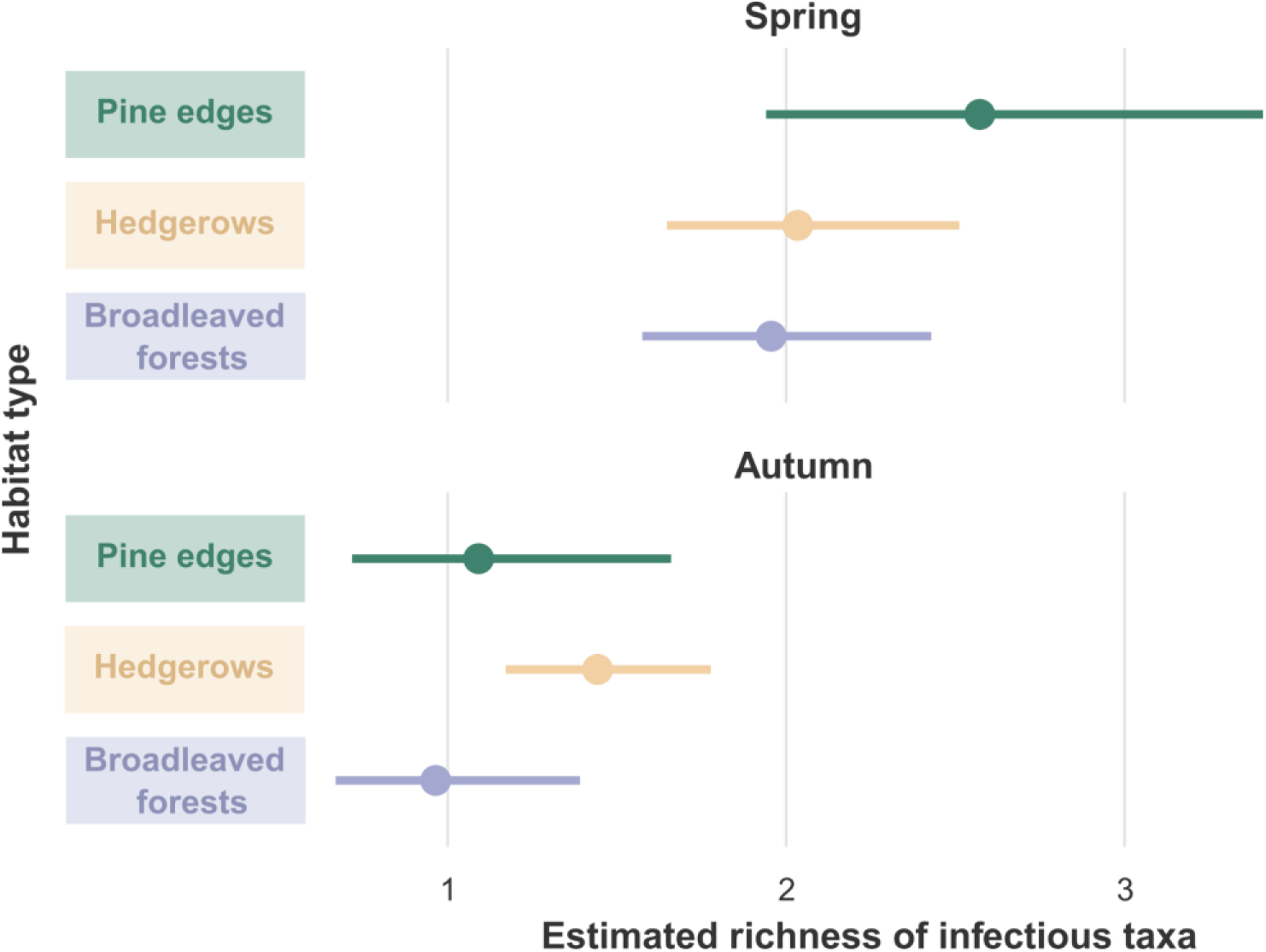
Estimated marginal means of infectious taxon richness across habitat types and seasons for *Apodemus sylvaticus* individuals. Bars represent 95% confidence intervals. Richness was significantly lower in Autumn (September-October) than in Spring (May-June) for all habitat categories (pine edge: *p* < 0.001; hedgerows: *p* = 0.011; broadleaved forest: *p* < 0.001). Habitat categories did not differ significantly within either season (Tukey-adjusted tests, all *p* > 0.05).

### 3.3 Infectious taxon-specific responses

#### 3.3.1 Infection probability

The best-supported HMSC model for infection probability included the habitat type, season, their interaction, local wood mouse abundance and host-level covariates (weight, sex, maturity). Across infectious taxa, this model achieved satisfactory MCMC convergence, with a mean explanatory power (Tjur’s R²) of 0.136 and a mean predictive power of 0.079 (HMSC_12_; β and Γ mean PSRFs < 1.002; Supplementary Table S4). Model performance varied across taxa with explanatory power ranging from 0.022 to 0.298 (Supplementary Table S5).

Weight accounted for the largest share of explained variation among fixed effects with 30.5% of the total explained variation on average followed by season (12.8%), habitat type (7.9%), habitat x season interaction (7%), local wood mouse abundance (4.3%), sex (3.6%) and maturity (3.5%). The remaining explained variation was distributed among random effects, including individual identity (9.8%), field mission (8.1%), spatial effect (6.6%), and site identity (5.9%).

Relative contribution of fixed effects varied among taxa, with season explaining larger variation for *Bartonella* spp. than for *Eperythrozoon* spp. (20.1% vs. 8% respectively, on average) for example, while weight accounted for a substantially greater share of explained variation in *Eperythrozoon* spp. than in *Bartonella* spp. (41.3% vs. 22.5% respectively; Figure 3B). Individual random effect contribution was also substantial for some taxa, notably *E. muris* (24.4%) and *E. coccoides* (17.9%).

**Figure 3.**
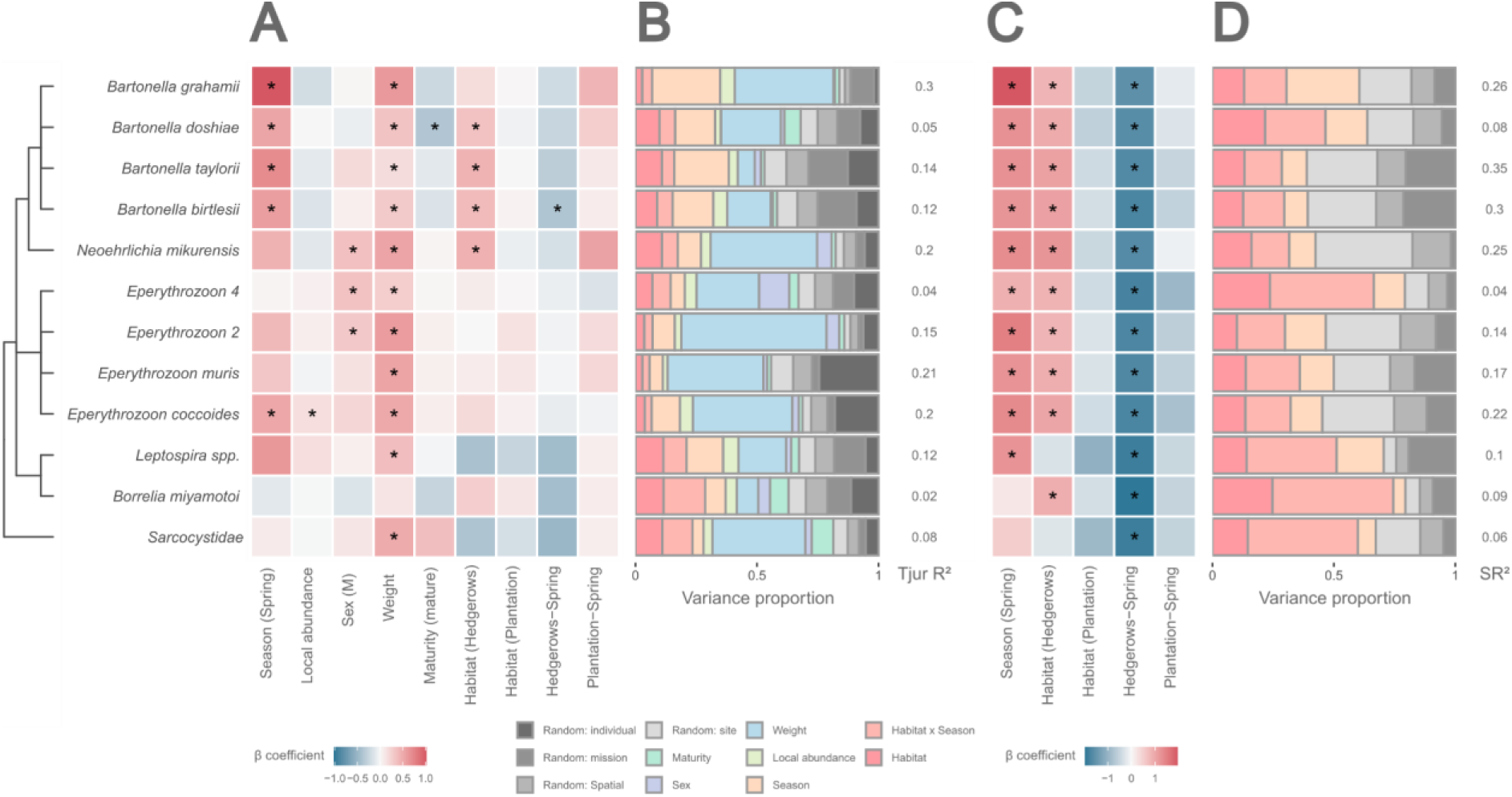
Posterior estimates of fixed effects (β parameters) and variance partitioning in the selected HMSC models. Results of the selected **A-B.** infection probability model (HMSC_12_) and **C-D.** infected-host abundance model (HMSC_17_). Both include habitat type, season and the interaction between season and habitat as fixed covariates with the addition of host characteristics for the infection probability model. The heatmaps **(A, C)** displays posterior mean effects of fixed covariates (β coefficients; columns) for each infectious taxon included in the analysis (rows). Asterisks indicate supported effects (PS ≥ 0.95), where PS is defined as the proportion of posterior samples sharing the same sign as the posterior mean. For categorical predictors, reference levels were broadleaved forest (habitat type), autumn (season), immature (host maturity), and female (host sex). The barplots **(B, D)** present the partitioning of explained variance among fixed covariates and random effects. For each infectious taxon (rows), bar heights indicate the relative contribution of each predictor to the explained variance. For each model, explanatory performance is indicated by Tjur’s R² and SR². Infectious taxa are ordered according to their phylogenetic relationships inferred from taxonomy. β coefficients are on model-specific scales and barplot colours consistently represent the same covariates across models.

Heavier individuals exhibited higher predicted infection probability for nearly all taxa, making host weight the most consistent host-level predictor (Figure 3A). In contrast, males showed higher predicted infection probabilities than females for three of the twelve infectious taxa, while maturity affected only *Bartonella doshiae*, for which mature individuals exhibited a lower infection probability. The probability of infection by *E. coccoides* was also positively impacted by local wood mouse abundance.

Across habitats, infection probability was either similar or higher in late spring than in autumn, with the largest and most consistent seasonal difference observed for *Bartonella grahamii* (Figure 4A).

**Figure 4.**
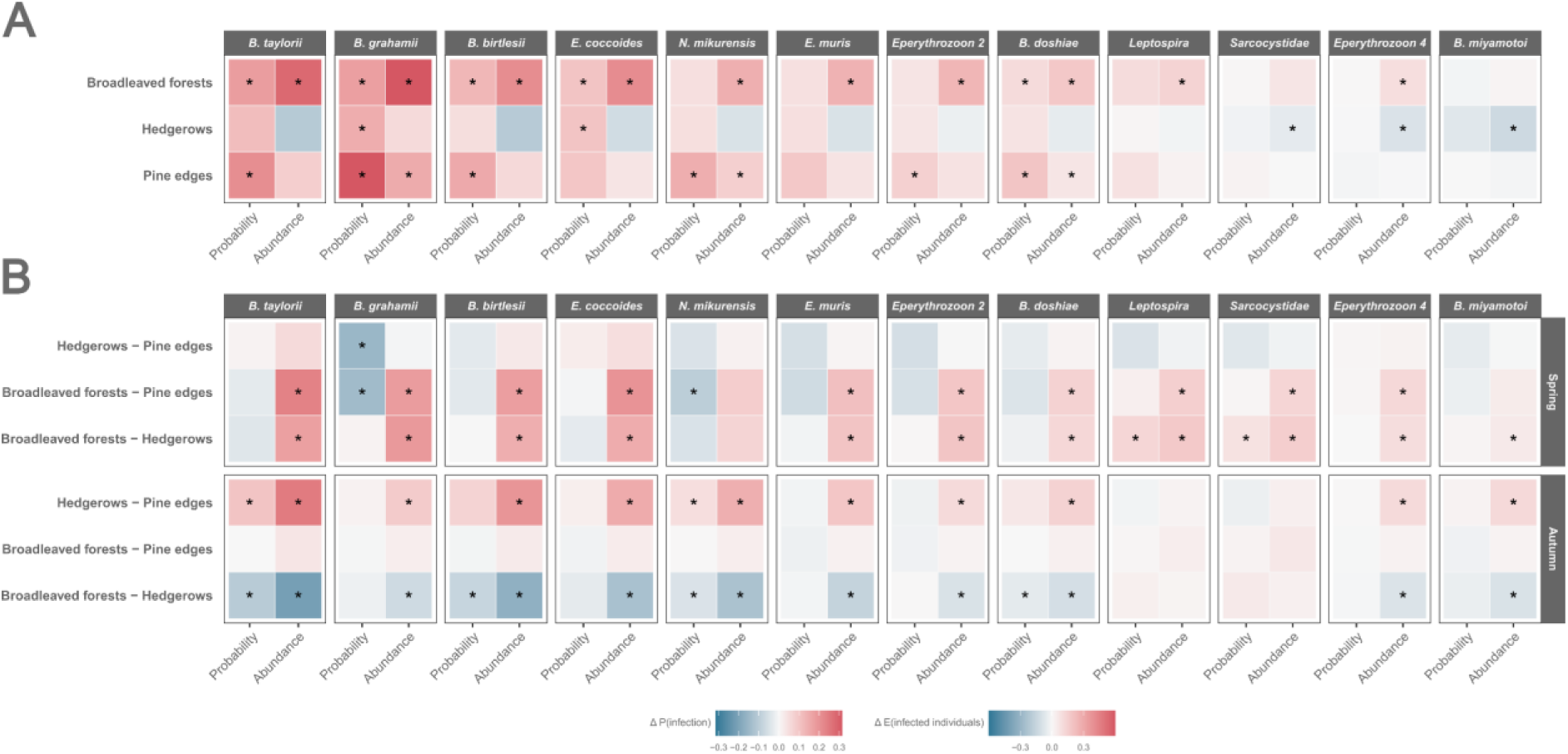
Seasonal and habitat effects on infection probability and infected-host abundance predicted by HMSC models. **A.** Pairwise differences in predicted marginal responses between spring and autumn (Spring - Autumn) for each habitat type (rows), infectious taxon (panels), and HMSC modelling approach (columns; Probability or Abundance). Responses correspond to either the predicted marginal infection probability (infection probability model) or the predicted marginal infected-host abundance (infected-host abundance model). **B.** Pairwise differences in predicted marginal responses for each pairwise habitat comparison (rows), season (row panels), infectious taxa (column panels), and HMSC approach (columns; Probability or Abundance). Responses correspond to either the predicted marginal infection probability (infection probability model) or the predicted marginal infected-host abundance (infected-host abundance model). Infectious taxa are ordered by decreasing mean effect size (average pairwise difference). Pairwise differences were calculated for each posterior sample as the difference between predicted marginal responses. Asterisks indicate supported contrasts (posterior support, PS ≥ 0.95), where PS is the proportion of posterior samples sharing the same sign as the posterior mean. Predicted differences are on different scales for the two modelling approaches.

In contrast, habitats’ effects differed markedly between seasons and infectious taxa (Figure 4B). During autumn, predicted infection probabilities were generally similar among habitats or tended to be higher in hedgerows than in the two other habitats, with supported or near-supported differences for *B. taylorii*, *B. birtlesii* and *N. mikurensis*. In spring, *B. grahamii* showed a higher predicted infection probability in pine edges than in hedgerows and broadleaved forests. Similarly, *N. mikurensis* exhibited a higher predicted infection probability in pine edges than in broadleaved forests. In contrast, *Leptospira* and *Sarcocystidae* spp. exhibited higher predicted infection probabilities in broadleaved forest than in hedgerows during this season.

Responses to fixed effects were more similar among phylogenetically close taxa (ρ = 0.926, Pr(ρ > 0) > 0.999). Vector-transmitted taxa tended to respond more positively to hedgerows relative to broadleaved forests than non-vector-transmitted taxa (Supplementary Figure S3). After accounting for fixed covariates and taxa traits, the residual variance-covariance matrix (Ω) provided no evidence of residual associations among infectious taxa (Supplementary Figure S4).

#### 3.3.2 Infected-host abundance

The best-supported HMSC model for infected-host abundance included the habitat type, season, and their interaction. Across infectious taxa, the selected model achieved satisfactory MCMC convergence, with a mean explanatory power (SR²) of 0.171 and a mean predictive power of 0.061 (HMSC_17_; β and Γ mean PSRFs < 1.019; Supplementary Table S4). Model explanatory performance varied across infectious taxa ranging from 0.042 to 0.345 (Supplementary Table S5).

The interaction between habitat and season accounted for the largest share of variation explained by fixed effects (27.1%), followed by habitat (15.9%) and season (13.9%). The remaining explained variation was distributed among random effects, including site identity (21.7%), field mission (10.9%), and spatial effect (10.5%; Figure 3D).

The relative contribution of model components varied among taxa (Figure 3B). For example, season accounted for a larger share of explained variation in *B. grahamii* (30%) than in other taxa, while site random effect accounted for a substantial share of variation in *N. mikurensis* (39.9%).

Predicted infected-host abundance was higher in late spring than in autumn except in hedgerows where infected-host abundance tended to be higher in autumn, especially for the least prevalent infectious taxa Sarcocystidae, *Eperythrozoon* 4 and *B. miyamotoi* (Figure 4A).

Habitat effects differed markedly between seasons but were consistent across infectious taxa within each season (Figure 4B). During autumn, predicted infected-host abundance was generally highest in hedgerows compared with the two other habitats. In contrast, during spring, broadleaved forests consistently exhibited the highest predicted infected-host abundance.

The abundance model showed an uncertain phylogenetic signal, with a bimodal posterior distribution (ρ = 0.717, Pr(ρ > 0) = 0.926). Vector-transmitted taxa tended to respond more positively to hedgerows relative to broadleaved forests than non-vector-transmitted taxa (Supplementary Figure S3). Supported residual associations were restricted to the session-level random effect (Supplementary Figure S4).

## 4. Discussion

In this study, we compared infection probability and infected-host abundance for a wide array of infectious taxa in *Apodemus sylvaticu*s, the only rodent species consistently persisting in the pine plantations of the Landes Forest, across three habitat types: pine plantation edges, broadleaved hedgerows, and natural broadleaved forests. We found that habitat effects on infection probability and infected-host abundance were contrasting and varied seasonally. Moreover habitat-related differences in host abundance generally outweighed those in infection probability in shaping hazard. Pine plantations generally supported fewer infected wood mice, whereas broadleaved hedgerows occasionally supported higher infected-host abundance than the two other habitats. Overall, our results provide insights into how forest plantation management can shape the infectiosome and zoonotic hazard through cascading effects on wildlife communities and pathogen circulation, while also showing that landscape restoration initiatives may have unexpected epidemiological consequences.

### 4.1 A diverse infectiosome including multiple zoonotic agents

Through the combination of multiple screening assays, we detected a broad diversity of infectious taxa in *Apodemus sylvaticus* from the Landes Forest, including nine zoonotic species known to cause human diseases. These findings further support the status of this widespread generalist rodent as a "hyper-reservoir" host (i.e., harboring more than three zoonotic pathogen species; Keesing and Ostfeld 2024). This role may be further enhanced by the frequent occurrence of co-infections, with almost half of individuals infected by multiple infectious agents. Our study also provided insights into the factors that may drive, or fail to drive, patterns of pathogen co-occurrence. Yet, we did not reveal any residual association between taxa at the individual level, while the few associations detected were linked to the field session, suggesting co-occurrences primarily related to temporal variations in wood mice abundance. The lack of such associations suggests either that interactions among co-occurring taxa are weak relative to shared environmental and host-related drivers, or that antagonistic interactions are counterbalanced by processes such as co-transmission (e.g. Moutailler et al., 2016) or shared exposure routes, or even that competition is reduced through diverse mechanisms such as fine-scale niche partitioning among these co-evolving organisms.

Notably, the infectious agents detected encompassed a wide range of taxa. No evidence of exposure to orthohantaviruses or orthoflaviviruses was detected. The absence of infection by orthohantaviruses is consistent with current knowledge, as no orthohantavirus has been described in *A. sylvaticus* from France yet. On the other hand, the mosquito-borne orthoflaviviruses Usutu virus (USUV) and West Nile virus (WNV) circulate endemically in southwestern France, as evidenced by reported infections in humans and wildlife during our sampling period (Duvignaud et al., 2024). To our knowledge, USUV has never been reported in *Apodemus spp.*, while WNV has only been detected once in *Apodemus flavicollis* (Cosseddu et al., 2017). Overall, the contribution of rodents to the ecology of these orthoflaviviruses remains poorly understood, and the few studies conducted in endemic areas have reported either no detection or low prevalence (Cosseddu et al., 2017; Diagne et al., 2019; Beaubaton et al., 2025). The absence of orthoflavivirus detection in our study should therefore not be interpreted as evidence that *Apodemus sylvaticus* is not exposed to these viruses, and continued surveillance is warranted.

Our detailed investigation of *Leptospira* diversity revealed the circulation of *L. interrogans* genovar Australis and *L. borgpetersenii*. Given that leptospirosis is one of the major rodent-borne zoonoses and has been a notifiable disease in humans in France since 2023 with a rising number of reported cases (Santé publique France, 2026), identifying the species and genovars circulating locally is of critical importance for public health. Because infection severity and host immunity against *Leptospira* is partially serovar-specific (Evangelista & Coburn, 2010), knowledge of local circulating strain is essential for guiding prevention strategies. These results therefore highlight the importance of wildlife surveillance not only for assessing local zoonotic risk but also for informing future vaccine development and adapting preventive measures for populations at risk of exposure.

Beyond the targeted screening assays, metabarcoding provided a broader picture of the infectiosome by simultaneously detecting multiple bacterial taxa without *a priori*, including zoonotic and non-zoonotic microorganisms. Most detected taxa were vector-borne hemotropic bacteria, mainly associated with lice, ticks and fleas like *Eperythrozoon* spp., *Borrelia* spp. and *Bartonella* spp.. This is expected, as *Apodemus* spp. play a major role in arthropod-borne transmission systems (Burri et al., 2014; Fabbri, Dorigo, et al., 2026; Perez et al., 2020). Among these bacteria, several neglected zoonotic agents were detected, including *Neoehrlichia mikurensis and Bartonella* spp., which are still scarcely targeted by routine species-specific assays (but see Michelet et al. 2014; Durand et al. 2026). *N. mikurensis* is a recently recognized tick-borne pathogen in Eurasia (Portillo et al., 2018), with increasing evidence pointing to *Apodemus* rodents as key reservoir hosts contributing to its maintenance in European enzootic cycles (Fabbri, Dorigo, et al., 2026). Together with the relatively high prevalence observed in our study, this supports the need to consider *N. mikurensis* in the differential diagnosis of post-tick bite febrile syndromes and reinforces the importance of prevention for forestry workers, hunters, and other professionals with high occupational exposure.

While most previous studies report *Bartonella* only at the genus level, or identify only the dominant sequence within samples through Sanger sequencing (Kosoy et al., 2018), our complementary metabarcoding approach provided species-level resolution while simultaneously detecting mixed infections. This revealed that nearly half infected individuals carried more than one *Bartonella* species, highlighting substantial within-host diversity. Although prevalence of *Bartonella* is high in many European rodents (Zarea et al., 2025), rodent-associated human bartonellosis remains rarely reported (Krügel et al., 2022).

While comprehensive, our characterization of the *A. sylvaticus* infectiosome nevertheless remains incomplete. Microorganisms exhibiting tissue tropism outside the bloodstream were probably underestimated by our spleen-based metabarcoding approach. For example, the zoonotic *Borrelia afzelii* is likely more reliably detected in skin than in spleen tissue (Zawada et al., 2020), suggesting that its prevalence, and consequently the associated zoonotic hazard, was underestimated in our study.

Some microorganisms detected incidentally warrant further investigation because of their potential zoonotic or veterinary importance. Organisms assigned to *Ehrlichia*, Sarcocystidae or *Streptobacillus* require additional characterization to determine their species identity and epidemiological relevance. *Streptobacillus,* in particular, has not previously been reported in *Apodemus sylvaticus*. This genus includes *Streptobacillus moniliformis*, the main causative agent of rat-bite fever, which is commonly associated with *Rattus norvegicus*, as well as several recently described *Streptobacillus* species associated with rat-bite fever-like illness in humans (Fukushima et al., 2018; Matt et al., 2021).

Besides the common rodent-associated hemoplasmas (i.e. *E. coccoides* and *E. muris*), several Mycoplasmotata OTUs could not be confidently assigned to a species, whereas others clustered with taxa initially described from humans or other vertebrates, including *Malacoplasma penetrans, Metamycoplasma sualvi* and *Mycoplasmopsis columbina.* These assignments may reflect the limited characterisation of some Mycoplasmatota (Gupta et al., 2018), with potentially undescribed rodent-associated lineages being assigned to their closest relatives rather than true species.

Lastly, additional groups of infectious agents, including helminths, unicellular eukaryotes and other virus families (e.g. poxviruses and coronaviruses), could be included to provide a wider picture of the wood mouse infectiosome.

### 4.2 Infectiosome dynamics are driven primarily by host characteristics and season

Host characteristics, particularly body mass, were the strongest predictors of individual variations in wood mouse infectiosome, outweighing environmental effects. Body mass was positively associated with infectious taxon richness, which is consistent with the accumulation of infections over lifetime. Because body mass is positively correlated with age in rodents and most of the infectious agents detected in this study are able to establish persistent infections, heavier individuals are likely to have had more time and opportunities to acquire infections.

Immature individuals also tended to be more frequently infected by *Bartonella* spp., with this pattern statistically supported for *B. doshiae*. These effects should be interpreted cautiously as maturity and body mass may capture different aspects of host demography and exposure history. In this case, higher infection probability for immature individuals might be explained by increased exposure to fleas during the period preceding their departure from the nest, when nest-associated flea transmission may be particularly important (Telfer, Begon, et al., 2007).

Sex also contributed to variation in infectiosome, with males showing higher infectious richness than females, a common pattern that may reflect sex-specific differences in ranging behaviour or immune investment (Klein, 2004; Zuk & McKean, 1996).

At the individual level, seasonality was the strongest environmental driver of the infectiosome and the second most important predictor overall after body mass, with infection probability consistently highest in spring. This pattern reflects a general feature of wildlife disease systems, where seasonal variation may emerge from the combined effects of environmentally induced changes in host behaviour, vector activity and host susceptibility (Altizer et al., 2006). In late spring, the onset of the breeding season may increase contact rates through increased movements and reproductive interactions (Attuquayefio et al., 1986; Wolton, 1985), while warmer temperatures may promote the activity of certain arthropod vectors such as questing ticks (Randolph, 2004), together increasing exposure opportunities. Simultaneously, hosts may be more susceptible during this period. Reproduction is energetically demanding and has been associated with reduced immune investment (Martin et al., 2007; Speakman, 2007). Late spring also marks a seasonal shift in the diet of wood mice, as declining seed availability leads to increased reliance on alternative food resources (Montgomery and Montgomery 1990), potentially increasing energetic constraints and further limiting investment in immunity (Forbes et al., 2016; Sweeny et al., 2021). The seasonal effects detected in our study were particularly strong for *Bartonella*. Such seasonal variation in *Bartonella* spp. infection has previously been reported in European rodents, with its direction and magnitude varying among *Bartonella* species. For example, *B. grahamii* showed particularly pronounced seasonal variation, with higher prevalence observed in autumn than in early spring (March), whereas *B. doshiae* showed the opposite trend (Telfer, Begon, et al., 2007; Telfer, Clough, et al., 2007). Here, *Bartonella* spp., and especially *B. grahamii*, consistently showed higher infection probability in late spring (May-June), when reproduction was already underway, than in autumn (September-October). *Bartonella* prevalence is influenced by both flea and rodent abundance (Telfer, Begon, et al., 2007), which may themselves exhibit different seasonal variations depending on the species and ecological systems (Hubbart et al., 2011; Huerta-Schliemann et al., 2025). These contrasting patterns may therefore reflect differences in local rodent communities and vector dynamics, as well as the timing of sampling relative to host reproduction.

Despite lower wood mouse abundance in spring than in autumn (Berard et al., 2026), the estimated number of infected individuals was overall higher in spring for most infectious taxa. This suggests that seasonal variation in infection probability can overshadow seasonal fluctuations in host abundance, potentially leading to an unexpected higher rodent-associated zoonotic hazard in spring. Hedgerows were an exception to this pattern. Although infection probability remained generally higher in spring for most taxa, the seasonal increase observed compared to autumn sessions was insufficient to offset the higher host abundance in autumn, resulting in a similar, or sometimes even greater, number of infected individuals for rare taxa (Sarcocystidae and *Leptospira* spp.) in autumn. These results highlight the importance of considering both host abundance and infection probability when assessing temporal and environmental variation in infection dynamics, as these measures do not necessarily vary in the same direction depending on the ecological systems studied. For example, de Cock et al. (2023) reported similar patterns for both measures in their study of wild rat infectiosome in urban settings, while Voutilainen et al. (2012) observed that lower infection probability could be associated with a higher abundance of Puumala virus-infected bank voles.

### 4.3 Pine plantation management may reduce infectious agent circulation through reduced wood mouse abundance

We expected pine plantation to increase infection probabilities, while their negative effect on host abundance was expected to result in variable patterns in the number of infected individuals across infectious taxa. In contrast, we found that pine edges had little consistent effect on infection probability and appeared to host less infected hosts for most infectious taxa. Besides, infectious taxon richness tended to be higher in pine plantations than in broadleaved forests, but in spring only. Moreover, higher infection probabilities were detected for only a few taxa, primarily the zoonotic agents *B. grahamii* and *N. mikurensis*. Nonetheless, these increases were consistently outweighed by the strong negative effect of pine plantations on host abundance, resulting in fewer infected wood mice in pine plantations compared to broadleaved forests for most infectious taxa.

Several non-mutually exclusive mechanisms could explain the higher infection probability observed for *B. grahamii* and *N. mikurensis* at pine edges. First, habitat-driven changes in host community composition may affect transmission patterns, depending on the relative abundance and competence of the hosts present. Changes in community composition may therefore either reduce transmission through the loss of competent hosts, generating a dilution effect, or increase it when competent hosts become proportionally dominant (Keesing et al., 2006; Keesing & Ostfeld, 2021). Compared to broadleaved forests, the pine plantation edges studied here support simplified vegetation resulting in lower small mammal diversity, particularly fewer bank voles (*Clethrionomys glareolus*; Berard et al., 2026). This pattern is likely to extend to other vertebrate groups, as structurally complex and diversified forests are associated with greater faunal diversity compared to tree monocultures (Ampoorter et al., 2020). Pine plantations may therefore promote community conditions less favourable to dilution for some infectious agents. Whether this mechanism contributes to the patterns observed for *B. grahamii* and *N. mikurensis* remains uncertain, however, as their host spectrum and host competence remain poorly characterised.

Second, habitat type may influence transmission by shaping arthropod vector communities. Vector assemblages are shaped by abiotic conditions, such as microclimate, which affect species-specific survival and activity (Krasnov et al., 2001; Leal et al., 2020), and by biotic factors such as the composition of vertebrate host communities, because ectoparasites differ in their host spectrum and specificity (Herrero-Cófreces et al., 2021; Krasnov et al., 2006). Habitat, season and their interaction may consequently generate distinct vector communities and phenologies (Krasnov et al., 1997; Herrero-Cófreces et al., 2021; Kahl & Gray, 2023; Benedek et al., 2024). If vector species differ in their abundance, competence or transmission efficiency, habitat-related shifts in their community composition and phenology could contribute to the season-specific infection patterns observed here.

Third, habitat-driven differences in individual susceptibility may also contribute to the observed patterns of infection probability. Although *A. sylvaticus* is considered an ecologically flexible species, populations do not perform equally well across habitat types (Unnsteinsdottir & Hersteinsson, 2011). Habitat characteristics, notably vegetation cover and composition, can influence perceived predation risk and food availability, thereby affecting chronic stress (Clinchy et al., 2013) and body condition (Parker et al., 2009; Vanroy et al., 2025), with potential cascading effects on immune function and infection susceptibility (Hing et al., 2016; Sánchez et al., 2018). Consistent with this, wood mice inhabiting structurally complex forests have been shown to exhibit better body condition (Vanroy et al., 2025), whereas individuals living in pine forests previously showed reduced body mass and elevated physiological stress (Navarro-Castilla & Barja, 2019). In our study, the lower resource availability and structural simplicity of pine plantations may therefore have exacerbated physiology constraints already experienced during spring, ultimately increasing host susceptibility to infection and leading to the season-specific pattern observed between pine edges and broadleaved forests. Further studies incorporating direct measures of physiological condition and immune function are needed to assess whether habitat-related physiological constraints contribute to variation in infection probability, particularly given that the observed pattern was detected for only two infectious taxa.

Nevertheless, these changes in infection probability remained too small to compensate for the much stronger effect of habitat on wood mouse abundance. Pine plantation edges supported substantially fewer wood mice than broadleaved forests, likely because their lower seed-productivity and simplified understorey provide fewer resources and less protective cover (Berard et al., 2026). Consequently, despite slightly higher infection probabilities for some taxa during spring, the estimated number of infected hosts was generally lower in pine plantations.

Overall, these results suggest that the effect of plantation management on infectious hazard may strongly depend on the ability of such habitats to sustain abundant populations of generalist hosts. Although forest plantations tend to filter out specialist species, the generalists that persist may not necessarily thrive either.

### 4.4 Broadleaved hedgerows generate distinct ecological conditions that shape the infectiosome in a taxon and season-dependent manner

Contrary to our expectation that broadleaved hedgerows would attenuate the epidemiological differences between pine plantation edges and broadleaved forests, hedgerows did not consistently exhibit an intermediate position along this habitat gradient. Instead, their effects were strongly season-dependent. In late spring, hedgerows showed some indications of attenuation relative to pine plantation edges, whereas in autumn they often exhibited higher infectious hazard than both other habitat types.

In late spring, several results were consistent with a limited attenuation of the epidemiological patterns observed at pine edges. Infectious taxon richness tended to be lower in hedgerows than at pine edges, and infection probability was lower for nine of the 12 infectious taxa (75%), although this difference was only statistically supported for *B. grahamii*. However, these differences in infection probability did not translate into a consistent reduction of infected-host abundance, which did not differ significantly between hedgerows and pine edges. Moreover, hedgerows did not consistently display the intermediate pattern expected between pine edges and broadleaved forests. The infectious taxon richness was similar in these habitats, as well as the probability of infection (except for two rare taxa, *Leptospira* spp. and Sarcocystidae). Instead, infected-host abundance was generally lower in hedgerows than in broadleaved forests for all infectious taxa except *N. mikurensis*.

This pattern was reversed in autumn. Hedgerows generally supported a higher infectious hazard than pine edges, with a trend toward greater infectious taxon richness and higher infected-host abundance for 10 of the 12 infectious taxa. Infection probabilities were also higher in hedgerows for *B. taylorii*, *B. birtlesii* and *N. mikurensis*. In parallel, hedgerows also exceeded broadleaved forests in infectious hazard. They tended to support greater infectious taxon richness, higher infection probabilities for these three taxa, and higher infected-host abundance for all but the same two rare taxa (*Leptospira* spp. and Sarcocystidae). Overall, these results suggest that hedgerows may exert little epidemiological effect beyond that observed at pine edges in late spring, but may elevate infectious hazard under autumnal conditions.

Several ecological and spatial characteristics of hedgerows may contribute to these contrasted seasonal results. As linear landscape elements, hedgerows can facilitate animal movements and concentrate activity (Davies & Pullin, 2007; Gelling et al., 2007), while also providing refuges when conditions in the surrounding matrix are less favourable (Tattersall et al., 2001). In our study system, these functions may become particularly important during autumnal acorn production, when oak-dominated hedgerows can provide concentrated food resources within an otherwise pine-dominated landscape. Higher food availability can reduce wood mouse home range size (Attuquayefio et al., 1986), potentially increasing local aggregation and contact rates among foraging individuals. At the same time, increased crowding may intensify competition and physiological stress, possibly leading to an increase in individual susceptibility to infection (Murray et al., 2016). In contrast, the lack of such a resource pulse in spring may reduce the ecological differences between hedgerows and pine edges, potentially contributing to their more similar epidemiological profiles observed during this season.

Microclimatic conditions, together with vertebrate host composition and abundance, may further shape vector assemblages differently across habitat types and seasons (Herrero-Cófreces et al., 2021; Benedek et al., 2024), potentially contributing to the season-dependent patterns in infectious hazard observed in these hedgerows. Compared with pine plantation edges, hedgerows harbour more diverse vegetation and fauna (Plat et al., 2026), while their linear configuration and their spatial extent distinguish them from broadleaved forests. In particular, although hedgerow vegetation may provide greater microclimatic buffering than pine edges, this effect is likely weaker than in broadleaved forests (Vanneste et al., 2020). These distinctive, environmental conditions may influence the composition and seasonal dynamics of vector communities, potentially favouring particular vector species involved in the transmission of wood mouse infectious taxa, or increasing vector-host encounters at specific times of the year. Assessing these processes could help explain why hedgerows appeared to provide more favourable conditions for transmission in autumn than in spring, and why their epidemiological profiles differed from those of both pine plantation edges and broadleaved forests.

### 4.5 Implications for forest management, ecological restoration and zoonotic surveillance

By comparing the infectiosome of *Apodemus sylvaticus* between natural broadleaved forests, hedgerows and pine plantations, our study provides new insights into how forest management and restoration can shape communities of infectious agents in wildlife, including zoonotic pathogens of public health concern.

Our findings first indicate that forest plantations can influence zoonotic hazard through their combined effects on infection probability and host abundance. Prevalence alone may provide a misleading picture of habitat-related changes in zoonotic hazard when habitat strongly influences host abundance, as seen here for pine plantations. Integrating both components provides a more informative assessment of how landscape management may alter pathogen persistence and transmission.

Our results also suggest that broadleaved forests may favour infectious taxa relying on environmental transmission. This was supported by the positive responses of *Leptospira* spp. and Sarcocystidae, as well as by the lower community-weighted mean of vector transmitted taxa observed in forests. The buffered microclimatic conditions and higher humidity prevailing in broadleaved forests may enhance the persistence of free-living infectious stages, such as Sarcocystidae oocysts (Lélu et al., 2012). In addition, the riparian broadleaved forests included in our study may provide additional opportunities for exposure to environmentally transmitted taxa. Water streams may accumulate and disperse free-living infectious stages, including leptospires washed into aquatic systems following rainfall events (Bierque et al., 2020).

Second, our results highlight that zoonotic hazards should be considered alongside biodiversity goals when planning or evaluating management interventions aimed at modifying forest composition and structure. This is especially relevant for forest diversification programmes or other restorative initiatives, including the establishment of broadleaved hedgerows, which are increasingly promoted to improve managed forest resilience and biodiversity in these forests (Messier et al., 2022).

Yet, our results show that potential restoration initiatives may sometimes result in a higher rodent-associated zoonotic hazard than both degraded and reference habitats. Indeed, broadleaved hedgerows did not consistently produce an epidemiological profile intermediate between pine plantations and broadleaved forests, and were associated with higher infectious hazard for several taxa in autumn. This should not be interpreted as an argument against ecological restoration, whose ecological benefits are well established (Atkinson et al., 2022; Benayas et al., 2009), but rather as a reminder that restoration interventions can generate context-dependent and sometimes unintended epidemiological consequences (Prist et al., 2023). This is especially important given the diversity of possible restoration approaches (Corlett, 2016) and the fact that most restored ecosystems do not end up recovering all characteristics of reference ecosystems (Atkinson et al., 2022).

Altogether, these findings argue for integrating disease ecology into the planning and evaluation of restoration initiatives to maximize ecological benefits while anticipating potential epidemiological consequences. Such an approach could help identify situations where appropriate, targeted surveillance and prevention efforts are most needed (Ecke et al., 2025).

### 4.6 Limitations and perspectives

Our estimates of rodent-associated zoonotic hazard were focused on *Apodemus sylvaticus*, which accounted for the vast majority of rodent captures across the study area. This provides a robust assessment of habitat-related patterns for this widespread generalist host, but does not capture the contribution of other host species. In particular, the bank vole (*Clethrionomys glareolus*), a recognised host of several zoonotic pathogens (Bouilloud et al., 2026), was captured in our study area, but almost exclusively in broadleaved forests (Berard et al., 2026). Its near absence from pine plantations prevented us from incorporating its contribution to pathogen circulation in our analyses. This may therefore have led to an incomplete assessment of differences in rodent-associated zoonotic hazard between habitats. Extending the comparison to the full small-mammal community would nevertheless be important to determine how habitat-driven changes in host community composition affect overall infectious hazard.

Rodent-associated zoonotic hazard should not be directly equated with human infection risk, which additionally depends on patterns of human exposure and preventive measures (e.g. Eisen and Dolan 2016). This distinction is particularly important for vector-borne pathogens, for which exposure involves several interacting compartments. For example, regarding tick-borne pathogens, the density of infected ticks is likely to be a more direct determinant of human exposure than the abundance of infected rodents (Gandy et al., 2022). Our results should therefore be interpreted as describing habitat-related changes in one wildlife component of zoonotic hazard, rather than changes in human infection risk.

A further limitation is that our comparison was based on mature broadleaved forests, mature hedgerows and mature pine plantations. As such, it provides a comparison among established habitat states rather than along a restoration or management trajectory. Small mammal communities are highly sensitive to forest succession stage and stand structure (Fuentes-Montemayor et al., 2020). Terrestrial rodent abundance may, for example, increase following clear-cutting because of the development of the understorey (Michał & Rafał, 2014), while hedgerow age also strongly determines vegetation structure (Montgomery et al. 2020). Therefore, zoonotic hazard is also likely to vary during plantation development or ecological restoration. Longitudinal studies following forest conversion or restoration would therefore be prioritized to identify when epidemiological changes occur and whether transient periods of increased hazard arise during these transitions (Prist et al., 2023).

In addition, forest plantations encompass highly diverse systems that differ widely in tree species, management intensity, landscape context and biogeographic conditions. The patterns observed in our study and the underlying ecological mechanisms proposed may therefore not operate similarly across plantation ecosystems and regions. Extending similar comparative studies across other contrasted plantation landscapes will therefore be essential to determine the generality of the habitat-driven epidemiological patterns observed here and to identify the conditions under which they emerge. Such comparisons would provide a stronger basis for designing restoration strategies that jointly consider biodiversity, ecosystem functioning and health.

## 5. Acknowledgements

We are grateful to the members of the Biodiversité Gènes et Communautés (INRAE-BIOGECO) team, particularly Nattan Plat, J-B. Rivoal, and Hervé Jactel, for their invaluable contributions to site selection, their help in facilitating site access, and for welcoming us to their laboratory. We also thank the management team of the Réserve Géologique de Saucats et La Brède National Nature Reserve (especially Yves Gilly) and the National Forests Office (ONF) for their support in granting permissions, offering guidance, and facilitating site access. We are also grateful to everyone who contributed to fieldwork during previous sampling campaigns, including Anne Loiseau, Sylvain Piry, Julie Blanchet, Karine Berthier, Florent Sebbane, and the Pasteur Lille team. We thank Loïc Pagès and Vincent Sluydts for valuable discussions and advice on joint species distribution modelling, and Benjamin Zeldvelder for his assistance with phylogenetic analyses. We would also like to acknowledge Audrey Weber (INRAE-AGAP) for the MiSeq sequencing and the GenoToul bioinformatics platform Toulouse Occitanie (GenoToul-Bioinfo, doi: 10.15454/1.5572369328961167E12) for providing support and computing resources.

## 6. Data, scripts, code, and supplementary information availability

The scripts and datasets supporting this study are available on GitHub (https://github.com/aoberard/Berard_infection_plantation).

## References

Ahlström, A., Canadell, J. G., & Metcalfe, D. B. (2022). Widespread Unquantified Conversion of Old Boreal Forests to Plantations. Earth’s Future, 10(11), e2022EF003221. 10.1029/2022EF003221

Altizer, S., Dobson, A., Hosseini, P., Hudson, P., Pascual, M., & Rohani, P. (2006). Seasonality and the dynamics of infectious diseases. Ecology Letters, 9(4), 467–484. 10.1111/j.1461-0248.2005.00879.x

Ampoorter, E., Barbaro, L., Jactel, H., Baeten, L., Boberg, J., Carnol, M., Castagneyrol, B., Charbonnier, Y., Dawud, S. M., Deconchat, M., Smedt, P. D., Wandeler, H. D., Guyot, V., Hättenschwiler, S., Joly, F.-X., Koricheva, J., Milligan, H., Muys, B., Nguyen, D.,… Allan, E. (2020). Tree diversity is key for promoting the diversity and abundance of forest-associated taxa in Europe. Oikos, 129(2), 133–146. 10.1111/oik.06290

Atkinson, J., Brudvig, L. A., Mallen-Cooper, M., Nakagawa, S., Moles, A. T., & Bonser, S. P. (2022). Terrestrial ecosystem restoration increases biodiversity and reduces its variability, but not to reference levels: A global meta-analysis. Ecology Letters, 25(7), 1725–1737. 10.1111/ele.14025

Attuquayefio, D. K., Gorman, M. L., & Wolton, R. J. (1986). Home range sizes in the Wood mouse Apodemus sylvaticus: Habitat, sex and seasonal differences. Journal of Zoology, 210(1), 45–53. 10.1111/j.1469-7998.1986.tb03619.x

Baker, R. E., Mahmud, A. S., Miller, I. F., Rajeev, M., Rasambainarivo, F., Rice, B. L., Takahashi, S., Tatem, A. J., Wagner, C. E., Wang, L.-F., Wesolowski, A., & Metcalf, C. J. E. (2022). Infectious disease in an era of global change. Nature Reviews Microbiology, 20(4), Article 4. 10.1038/s41579-021-00639-z

Beaubaton, R., Revel, J., Pigeyre, L., Bollore, K., Lepeule, A., Mocq, J., Franceschi, C. de, Pradel, J., Perrin, Y., Gomis, D., Ducousso, M., Virolle, L., Chenet, B., Castel, G., Charmantier, A., Charbonnel, N., Lacour, G., Courot, O., Mignotte, A., & Simonin, Y. (2025). Tracking the urban spread of Usutu virus in southern France: Detection across biological and environmental matrices. PLOS Neglected Tropical Diseases, 19(9), e0013506. 10.1371/journal.pntd.0013506

Benayas, J. M. R., Newton, A. C., Diaz, A., & Bullock, J. M. (2009). Enhancement of Biodiversity and Ecosystem Services by Ecological Restoration: A Meta-Analysis. Science, 325(5944), 1121–1124. 10.1126/science.1172460

Benedek, A. M., Boeraș, I., Lazăr, A., Sandu, A., Cocîrlea, M. D., Stănciugelu, M., Cic, N. V., & Postolache, C. (2024). Effects of Season, Habitat, and Host Characteristics on Ectoparasites of Wild Rodents in a Mosaic Rural Landscape. Animals, 14(2), 304. 10.3390/ani14020304

Berard, A., Plat, N., Pradel, J., Galan, M., Loiseau, A., Piry, S., Blanchet, J., Cesari, L., Berthier, K., Rivoal, J.-B., Pellett, C., Valbuena, R., Jactel, H., & Charbonnel, C. (2026). Broadleaved hedgerows as complementary habitats for small mammals in pine plantation forest landscapes. Forest Ecology and Management, 621, 124259. 10.1016/j.foreco.2026.124259

Bierque, E., Thibeaux, R., Girault, D., Soupé-Gilbert, M.-E., & Goarant, C. (2020). A systematic review of Leptospira in water and soil environments. PLOS ONE, 15(1), e0227055. 10.1371/journal.pone.0227055

Bouilloud, M., Galan, M., Bordes, A., Tatard, C., Gauthier, P., Pradel, J., Castel, G., Alburkat, H., Dutra, L., Sironen, T., Galon, C., Moutailler, S., Djelouadji, Z., Roche, B., & Charbonnel, N. (2026). Surveillance of Zoonotic Pathogens in Small Mammals Across Forests With Different Levels of Anthropization in Eastern France. Transboundary and Emerging Diseases, 2026(1), 4038648. 10.1155/tbed/4038648

Brockerhoff, E. G., Barbaro, L., Castagneyrol, B., Forrester, D. I., Gardiner, B., González-Olabarria, J. R., Lyver, P. O., Meurisse, N., Oxbrough, A., Taki, H., Thompson, I. D., van der Plas, F., & Jactel, H. (2017). Forest biodiversity, ecosystem functioning and the provision of ecosystem services. Biodiversity and Conservation, 26(13), 3005–3035. 10.1007/s10531-017-1453-2

Brockerhoff, E. G., Jactel, H., Parrotta, J. A., Quine, C. P., & Sayer, J. (2008). Plantation forests and biodiversity: Oxymoron or opportunity? Biodiversity and Conservation, 17(5), 925–951. 10.1007/s10531-008-9380-x

Bugarski-Stanojević, V., Blagojević, J., Adna?ević, T., Jojić, V., & Vujošević, M. (2008). Molecular phylogeny and distribution of three Apodemus species (Muridae, Rodentia) in Serbia. Journal of Zoological Systematics and Evolutionary Research, 46(3), 278–286. 10.1111/j.1439-0469.2007.00452.x

Burri, C., Schumann, O., Schumann, C., & Gern, L. (2014). Are Apodemus spp. Mice and Myodes glareolus reservoirs for Borrelia miyamotoi, Candidatus Neoehrlichia mikurensis, Rickettsia helvetica, R. monacensis and Anaplasma phagocytophilum? Ticks and Tick-Borne Diseases, 5(3), 245–251. 10.1016/j.ttbdis.2013.11.007

Carlson, C. J., Brookson, C. B., Becker, D. J., Cummings, C. A., Gibb, R., Halliday, F. W., Heckley, A. M., Huang, Z. Y. X., Lavelle, T., Robertson, H., Vicente-Santos, A., Weets, C. M., & Poisot, T. (2025). Pathogens and planetary change. Nature Reviews. Biodiversity, 1(1), 32–49. 10.1038/s44358-024-00005-w

Chaudhary, A., Burivalova, Z., Koh, L. P., & Hellweg, S. (2016). Impact of Forest Management on Species Richness: Global Meta-Analysis and Economic Trade-Offs. Scientific Reports, 6(1), 23954. 10.1038/srep23954

Clinchy, M., Sheriff, M. J., & Zanette, L. Y. (2013). Predator-induced stress and the ecology of fear. Functional Ecology, 27(1), 56–65. 10.1111/1365-2435.12007

Corlett, R. T. (2016). Restoration, Reintroduction, and Rewilding in a Changing World. Trends in Ecology & Evolution, 31(6), 453–462. 10.1016/j.tree.2016.02.017

Cosseddu, G. M., Sozio, G., Valleriani, F., Di Gennaro, A., Pascucci, I., Gavaudan, S., Marianneau, P., & Monaco, F. (2017). Serological Survey of Hantavirus and Flavivirus Among Wild Rodents in Central Italy. Vector-Borne and Zoonotic Diseases, 17(11), 777–779. 10.1089/vbz.2017.2143

Davidson, G., Speldewinde, P., Manin, B. O., Cook, A., Weinstein, P., & Chua, T. H. (2024). Forest Restoration and the Zoonotic Vector Anopheles balabacensis in Sabah, Malaysia. EcoHealth, 21(1), 21–37. 10.1007/s10393-024-01675-w

Davies, Z. G., & Pullin, A. S. (2007). Are hedgerows effective corridors between fragments of woodland habitat? An evidence-based approach. Landscape Ecology, 22(3), 333–351. 10.1007/s10980-006-9064-4

de Cock, M. P., de Vries, A., Fonville, M., Esser, H. J., Mehl, C., Ulrich, R. G., Joeres, M., Hoffmann, D., Eisenberg, T., Schmidt, K., Hulst, M., van der Poel, W. H. M., Sprong, H., & Maas, M. (2023). Increased rat-borne zoonotic disease hazard in greener urban areas. Science of The Total Environment, 896, 165069. 10.1016/j.scitotenv.2023.165069

Diagne, M. M., Ndione, M. H. D., Di Paola, N., Fall, G., Bedekelabou, A. P., Sembène, P. M., Faye, O., Zanotto, P. M. de A., & Sall, A. A. (2019). Usutu Virus Isolated from Rodents in Senegal. Viruses, 11(2), 181. 10.3390/v11020181

Dobigny, G., Garba, M., Tatard, C., Loiseau, A., Galan, M., Kadaouré, I., Rossi, J.-P., Picardeau, M., & Bertherat, E. (2015). Urban Market Gardening and Rodent-Borne Pathogenic Leptospira in Arid Zones: A Case Study in Niamey, Niger. PLoS Neglected Tropical Diseases, 9(10), e0004097. 10.1371/journal.pntd.0004097

Dubois, A., Castel, G., Murri, S., Pulido, C., Pons, J.-B., Benoit, L., Loiseau, A., Lakhdar, L., Galan, M., Charbonnel, N., & Marianneau, P. (2017). Experimental infections of wild bank voles (Myodes glareolus) from nephropatia epidemica endemic and non-endemic regions revealed slight differences in Puumala virological course and immunological responses. Virus Research, 235, 67–72. 10.1016/j.virusres.2017.04.004

Durand, J., Bah, T.-M., Lebert, I., Galon, C., Carravieri, I., Masseglia, S., Armand, J.-M., Marchand, J., Galley, C., Chalvet-Monfray, K., Vayssier-Taussat, M., Vourc’h, G., Brun-Jacob, A., Moutailler, S., Bailly, X., & Frey-Klett, P. (2026). Distribution of tick-borne microorganisms in human-biting ticks in France collected through a Citizen-science program. Ticks and Tick-Borne Diseases, 17(2), 102612. 10.1016/j.ttbdis.2026.102612

Duvignaud, A., Bigeard, C., Fontaine, A., Grard, G., Touzet, T., Migne, C., Gallian, P., Malvy, D., de Lamballerie, X., & Gonzalez, G. (2024). Investigation One Health de l’émergence des virus West Nile et Usutu en Nouvelle Aquitaine. Médecine et Maladies Infectieuses Formation, 24es Journées Nationales d’Infectiologie, 3(2, Supplement), S9. 10.1016/j.mmifmc.2024.04.018

Ecke, F., Semenza, J. C., Buzan, E., Costa, F., Giorgi, E., Guo, J., Kirkpatrick, L., Knauf, S., Meheretu, Y., Singh, N., Sjödin, H., Timperley, M., Treskova, M., Ulrich, R. G., Zeppelini, C. G., & Rocklöv, J. (2025). Adaptive ecosystem restoration to mitigate zoonotic risks. Nature Ecology & Evolution, 9(11), 1979–1988. 10.1038/s41559-025-02869-3

Eisen, L., & Dolan, M. C. (2016). Evidence for Personal Protective Measures to Reduce Human Contact With Blacklegged Ticks and for Environmentally Based Control Methods to Suppress Host-Seeking Blacklegged Ticks and Reduce Infection with Lyme Disease Spirochetes in Tick Vectors and Rodent Reservoirs. Journal of Medical Entomology, tjw103. 10.1093/jme/tjw103

Escudié, F., Auer, L., Bernard, M., Mariadassou, M., Cauquil, L., Vidal, K., Maman, S., Hernandez-Raquet, G., Combes, S., & Pascal, G. (2018). FROGS: Find, Rapidly, OTUs with Galaxy Solution. Bioinformatics, 34(8), 1287–1294. 10.1093/bioinformatics/btx791

Evangelista, K. V., & Coburn, J. (2010). Leptospira as an emerging pathogen: A review of its biology, pathogenesis and host immune responses. Future Microbiology, 5(9), 1413–1425. 10.2217/fmb.10.102

Fabbri, D., Dorigo, L., Grassi, L., Leopardi, S., Tagliapietra, V., & Beraldo, P. (2026). From reservoirs to ecological integrators: The role of European Apodemus spp. in vector-borne zoonotic pathogens. Frontiers in Veterinary Science, 13. 10.3389/fvets.2026.1822333

Fabbri, D., Grassi, L., Dorigo, L., Leopardi, S., Tagliapietra, V., & Beraldo, P. (2026). Small rodents, huge impact: A scoping review of European *Apodemus* species in the ecology of directly transmitted, environmentally mediated, and interface-associated zoonotic pathogens. One Health, 23, 101532. 10.1016/j.onehlt.2026.101532

FAO. (2025). Global Forest Resources Assessment 2025. FAO. 10.4060/cd6709en

Fell, A., Jagadesh, S., Duthie, A. B., Kirkpatrick, L., & Bunnefeld, N. (2026). Global evidence synthesis on land-use change and zoonotic risks. Nature Sustainability, 9(1), 142–152. 10.1038/s41893-025-01750-2

Fisher, M. C., & Garner, T. W. J. (2020). Chytrid fungi and global amphibian declines. Nature Reviews Microbiology, 18(6), 332–343. 10.1038/s41579-020-0335-x

Forbes, K. M., Mappes, T., Sironen, T., Strandin, T., Stuart, P., Meri, S., Vapalahti, O., Henttonen, H., & Huitu, O. (2016). Food limitation constrains host immune responses to nematode infections. Biology Letters, 12(9), 20160471. 10.1098/rsbl.2016.0471

Fuentes-Montemayor, E., Ferryman, M., Watts, K., Macgregor, N. A., Hambly, N., Brennan, S., Coxon, R., Langridge, H., & Park, K. J. (2020). Small mammal responses to long-term large-scale woodland creation: The influence of local and landscape-level attributes. Ecological Applications, 30(2), e02028. 10.1002/eap.2028

Fukushima, K., Yanagisawa, N., Imaoka, K., Kimura, M., & Imamura, A. (2018). Rat-bite fever due to *Streptobacillus notomytis* isolated from a human specimen. Journal of Infection and Chemotherapy, 24(4), 302–304. 10.1016/j.jiac.2017.10.018

Galan, M., Pons, J.-B., Tournayre, O., Pierre, É., Leuchtmann, M., Pontier, D., & Charbonnel, N. (2018). Metabarcoding for the parallel identification of several hundred predators and their prey: Application to bat species diet analysis. Molecular Ecology Resources, 18(3), 474–489. 10.1111/1755-0998.12749

Galan, M., Razzauti, M., Bard, E., Bernard, M., Brouat, C., Charbonnel, N., Dehne-Garcia, A., Loiseau, A., Tatard, C., Tamisier, L., Vayssier-Taussat, M., Vignes, H., & Cosson, J.-F. (2016). 16S rRNA Amplicon Sequencing for Epidemiological Surveys of Bacteria in Wildlife. mSystems, 1(4), e00032–16. 10.1128/mSystems.00032-16

Gandy, S., Kilbride, E., Biek, R., Millins, C., & Gilbert, L. (2022). No net effect of host density on tick-borne disease hazard due to opposing roles of vector amplification and pathogen dilution. Ecology and Evolution, 12(9), e9253. 10.1002/ece3.9253

Garcia-Lopez, M., Lurier, T., Bouilloud, M., Pradel, J., Tatard, C., Sepulveda, D., Anfray, G., Dussert, J., Bourhy, P., Charbonnel, N., & Djelouadji, Z. (2024). Prevalence, genetic diversity and eco-epidemiology of pathogenic Leptospira species in small mammal communities in urban parks Lyon city, France. PLOS ONE, 19(4), e0300523. 10.1371/journal.pone.0300523

Gelling, M., Macdonald, D. W., & Mathews, F. (2007). Are hedgerows the route to increased farmland small mammal density? Use of hedgerows in British pastoral habitats. Landscape Ecology, 22(7), 1019–1032. 10.1007/s10980-007-9088-4

Gelman, A., & Rubin, D. B. (1992). Inference from Iterative Simulation Using Multiple Sequences. Statistical Science, 7(4), 457–472. 10.1214/ss/1177011136

Gibb, R., Redding, D. W., Chin, K. Q., Donnelly, C. A., Blackburn, T. M., Newbold, T., & Jones, K. E. (2020). Zoonotic host diversity increases in human-dominated ecosystems. Nature, 584(7821), Article 7821. 10.1038/s41586-020-2562-8

Gottdenker, N. L., Streicker, D. G., Faust, C. L., & Carroll, C. R. (2014). Anthropogenic Land Use Change and Infectious Diseases: A Review of the Evidence. EcoHealth, 11(4), 619–632. 10.1007/s10393-014-0941-z

Guégan, J.-F., Ayouba, A., Cappelle, J., & de Thoisy, B. (2020). Forests and emerging infectious diseases: Unleashing the beast within. Environmental Research Letters, 15(8), 083007. 10.1088/1748-9326/ab8dd7

Guégan, J.-F., de Thoisy, B., Gomez-Gallego, M., & Jactel, H. (2023). World forests, global change, and emerging pests and pathogens. Current Opinion in Environmental Sustainability, 61, 101266. 10.1016/j.cosust.2023.101266

Gupta, R. S., Sawnani, S., Adeolu, M., Alnajar, S., & Oren, A. (2018). Phylogenetic framework for the phylum Tenericutes based on genome sequence data: Proposal for the creation of a new order Mycoplasmoidales ord. nov., containing two new families Mycoplasmoidaceae fam. nov. and Metamycoplasmataceae fam. nov. harbouring Eperythrozoon, Ureaplasma and five novel genera. Antonie van Leeuwenhoek, 111(9), 1583–1630. 10.1007/s10482-018-1047-3

Han, B. A., Schmidt, J. P., Bowden, S. E., & Drake, J. M. (2015). Rodent reservoirs of future zoonotic diseases. Proceedings of the National Academy of Sciences, 112(22), 7039–7044. 10.1073/pnas.1501598112

Heni, A. C., Fackelmann, G., Eibner, G., Kreinert, S., Schmid, J., Schwensow, N. I., Wiegand, J., Wilhelm, K., & Sommer, S. (2023). Wildlife gut microbiomes of sympatric generalist species respond differently to anthropogenic landscape disturbances. Animal Microbiome, 5(1), 22. 10.1186/s42523-023-00237-9

Herrero-Cófreces, S., Flechoso, M. F., Rodríguez-Pastor, R., Luque-Larena, J. J., & Mougeot, F. (2021). Patterns of flea infestation in rodents and insectivores from intensified agro-ecosystems, Northwest Spain. Parasites & Vectors, 14(1), 16. 10.1186/s13071-020-04492-6

Hing, S., Narayan, E. J., Thompson, R. C. A., & Godfrey, S. S. (2016). The relationship between physiological stress and wildlife disease: Consequences for health and conservation. Wildlife Research, 43(1), 51–60. 10.1071/WR15183

Hoyt, J. R., Kilpatrick, A. M., & Langwig, K. E. (2021). Ecology and impacts of white-nose syndrome on bats. Nature Reviews Microbiology, 19(3), 196–210. 10.1038/s41579-020-00493-5

Hua, F., Wang, L., Fisher, B., Zheng, X., Wang, X., Yu, D. W., Tang, Y., Zhu, J., & Wilcove, D. S. (2018). Tree plantations displacing native forests: The nature and drivers of apparent forest recovery on former croplands in Southwestern China from 2000 to 2015. Biological Conservation, 222, 113–124. 10.1016/j.biocon.2018.03.034

Hubbart, J. A., Jachowski, D. S., & Eads, D. A. (2011). Seasonal and among-site variation in the occurrence and abundance of fleas on California ground squirrels (Otospermophilus beecheyi). Journal of Vector Ecology: Journal of the Society for Vector Ecology, 36(1), 117–123. 10.1111/j.1948-7134.2011.00148.x

Huerta-Schliemann, L. de la, Vilella, M., Freixas, L., & Torre, I. (2025). Effects of Climate and Land Use on the Population Dynamics of the Bank Vole (Clethrionomys glareolus) in the Southernmost Part of Its Range. Animals, 15(6). 10.3390/ani15060839

Iezzi, M. E., Cruz, P., Varela, D., De Angelo, C., & Di Bitetti, M. S. (2018). Tree monocultures in a biodiversity hotspot: Impact of pine plantations on mammal and bird assemblages in the Atlantic Forest. Forest Ecology and Management, 424, 216–227. 10.1016/j.foreco.2018.04.049

Infante, J., Riquelme, M., Huerta, N., Oettinger, S., Fredes, F., Simonetti, J. A., & Rubio, A. V. (2022). *Cryptosporidium* spp. and *Giardia* spp. in wild rodents: Using occupancy models to estimate drivers of occurrence and prevalence in native forest and exotic *Pinus radiata* plantations from Central Chile. Acta Tropica, 235, 106635. 10.1016/j.actatropica.2022.106635

Jolivet, C., Augusto, L., Trichet, P., & Arrouays, D. (2007). Les sols du massif forestier des Landes de Gascogne: Formation, histoire, propriétés et variabilité spatiale. Revue Forestière Française, 59(1), 7–30. 10.4267/2042/8480

Jones, K. E., Patel, N. G., Levy, M. A., Storeygard, A., Balk, D., Gittleman, J. L., & Daszak, P. (2008). Global trends in emerging infectious diseases. Nature, 451(7181), 990–993. 10.1038/nature06536

Kahl, O., & Gray, J. S. (2023). The biology of *Ixodes ricinus* with emphasis on its ecology. Ticks and Tick-Borne Diseases, 14(2), 102114. 10.1016/j.ttbdis.2022.102114

Keesing, F., Holt, R. D., & Ostfeld, R. S. (2006). Effects of species diversity on disease risk. Ecology Letters, 9(4), 485–498. 10.1111/j.1461-0248.2006.00885.x

Keesing, F., & Ostfeld, R. S. (2021). Dilution effects in disease ecology. Ecology Letters, 24(11), 2490–2505. 10.1111/ele.13875

Keesing, F., & Ostfeld, R. S. (2024). Emerging patterns in rodent-borne zoonotic diseases. Science, 385(6715), 1305–1310. 10.1126/science.adq7993

Klein, S. L. (2004). Hormonal and immunological mechanisms mediating sex differences in parasite infection. Parasite Immunology, 26(6–7), 247–264. 10.1111/j.0141-9838.2004.00710.x

Kosoy, M., Mckee, C., Albayrak, L., & Fofanov, Y. (2018). Genotyping of Bartonella bacteria and their animal hosts: Current status and perspectives. Parasitology, 145(5), 543–562. 10.1017/S0031182017001263

Krasnov, B. R., Khokhlova, I. S., Fielden, L. J., & Burdelova, N. V. (2001). Effect of Air Temperature and Humidity on the Survival of Pre-Imaginal Stages of Two Flea Species (Siphonaptera: Pulicidae). Journal of Medical Entomology, 38(5), 629–637. 10.1603/0022-2585-38.5.629

Krasnov, B. R., Shenbrot, G. I., Medvedev, S. G., Vatschenok, V. S., & Khokhlova, I. S. (1997). Host–habitat relations as an important determinant of spatial distribution of flea assemblages (Siphonaptera) on rodents in the Negev Desert. Parasitology, 114(2), 159–173. 10.1017/S0031182096008347

Krasnov, B. R., Stanko, M., Miklisova, D., & Morand, S. (2006). Habitat variation in species composition of flea assemblages on small mammals in central Europe. Ecological Research, 21(3), 460–469. 10.1007/s11284-005-0142-x

Krügel, M., Król, N., Kempf, V. A. J., Pfeffer, M., & Obiegala, A. (2022). Emerging rodent-associated Bartonella: A threat for human health? Parasites & Vectors, 15(1), 113. 10.1186/s13071-022-05162-5

Leal, B., Zamora, E., Fuentes, A., Thomas, D. B., & Dearth, R. K. (2020). Questing by Tick Larvae (Acari: Ixodidae): A Review of the Influences That Affect Off-Host Survival. Annals of the Entomological Society of America, 113(6), 425–438. 10.1093/aesa/saaa013

Lélu, M., Villena, I., Dardé, M.-L., Aubert, D., Geers, R., Dupuis, E., Marnef, F., Poulle, M.-L., Gotteland, C., Dumètre, A., & Gilot-Fromont, E. (2012). Quantitative Estimation of the Viability of Toxoplasma gondii Oocysts in Soil. Applied and Environmental Microbiology, 78(15), 5127–5132. 10.1128/AEM.00246-12

Martin, L. B., Weil, Z. M., & Nelson, R. J. (2007). Seasonal changes in vertebrate immune activity: Mediation by physiological trade-offs. Philosophical Transactions of the Royal Society B: Biological Sciences, 363(1490), 321–339. 10.1098/rstb.2007.2142

Matt, U., Schmiedel, J., Fawzy, A., Trauth, J., Schmidt, K., Vogel, K., Herold, S., Karrasch, T., Imirzalioglu, C., & Eisenberg, T. (2021). Infection in a Young Immunocompetent Male Caused by Streptobacillus felis, a Putative Zoonotic Microorganism Transmitted by Cats. Clinical Infectious Diseases, 72(10), 1826–1829. 10.1093/cid/ciaa968

Meadows, A. J., Stephenson, N., Madhav, N. K., & Oppenheim, B. (2023). Historical trends demonstrate a pattern of increasingly frequent and severe spillover events of high-consequence zoonotic viruses. BMJ Global Health, 8(11), e012026. 10.1136/bmjgh-2023-012026

Messier, C., Bauhus, J., Sousa-Silva, R., Auge, H., Baeten, L., Barsoum, N., Bruelheide, H., Caldwell, B., Cavender-Bares, J., Dhiedt, E., Eisenhauer, N., Ganade, G., Gravel, D., Guillemot, J., Hall, J. S., Hector, A., Hérault, B., Jactel, H., Koricheva, J.,… Zemp, D. C. (2022). For the sake of resilience and multifunctionality, let’s diversify planted forests! Conservation Letters, 15(1), e12829. 10.1111/conl.12829

Messina, S., Edwards, D. P., Eens, M., & Costantini, D. (2018). Physiological and immunological responses of birds and mammals to forest degradation: A meta-analysis. Biological Conservation, 224, 223–229. 10.1016/j.biocon.2018.06.002

Michał, B., & Rafał, Z. (2014). Responses of small mammals to clear-cutting in temperate and boreal forests of Europe: A meta-analysis and review. European Journal of Forest Research, 133(1), 1–11. 10.1007/s10342-013-0726-x

Michelet, L., Delannoy, S., Devillers, E., Umhang, G., Aspan, A., Juremalm, M., Chirico, J., van der Wal, F. J., Sprong, H., Boye Pihl, T. P., Klitgaard, K., Bødker, R., Fach, P., & Moutailler, S. (2014). High-throughput screening of tick-borne pathogens in Europe. Frontiers in Cellular and Infection Microbiology, 4. 10.3389/fcimb.2014.00103

Mills, J. N., Yates, T. L., Childs, J. E., Parmenter, R. R., Ksiazek, T. G., Rollin, P. E., & Peters, C. J. (1995). Guidelines for Working with Rodents Potentially Infected with Hantavirus. Journal of Mammalogy, 76(3), 716–722. 10.2307/1382742

Mollentze, N., & Streicker, D. G. (2020). Viral zoonotic risk is homogenous among taxonomic orders of mammalian and avian reservoir hosts. Proceedings of the National Academy of Sciences, 117(17), 9423–9430. 10.1073/pnas.1919176117

Montgomery, I., Caruso, T., & Reid, N. (2020). Hedgerows as Ecosystems: Service Delivery, Management, and Restoration. Annual Review of Ecology, Evolution, and Systematics, 51, 81–102. 10.1146/annurev-ecolsys-012120-100346

Montgomery, S. S. J., & Montgomery, W. I. (1990). Intrapopulation variation in the diet of the wood mouse Apodemus sylvaticus. Journal of Zoology, 222(4), 641–651. 10.1111/j.1469-7998.1990.tb06020.x

Mora, O., Banos, V., Regolini, M., & Carnus, J.-M. (2012). Le massif des Landes de Gascogne à l’horizon 2050. Rapport de l’étude prospective (p. 290). Conseil régional d’Aquitaine - INRA.

Morand, S., & Lajaunie, C. (2021). Outbreaks of Vector-Borne and Zoonotic Diseases Are Associated With Changes in Forest Cover and Oil Palm Expansion at Global Scale. Frontiers in Veterinary Science, 8. 10.3389/fvets.2021.661063

Moutailler, S., Moro, C. V., Vaumourin, E., Michelet, L., Tran, F. H., Devillers, E., Cosson, J.-F., Gasqui, P., Van, V. T., Mavingui, P., Vourc’h, G., & Vayssier-Taussat, M. (2016). Co-infection of Ticks: The Rule Rather Than the Exception. PLOS Neglected Tropical Diseases, 10(3), e0004539. 10.1371/journal.pntd.0004539

Murray, M. H., Becker, D. J., Hall, R. J., & Hernandez, S. M. (2016). Wildlife health and supplemental feeding: A review and management recommendations. Biological Conservation, 204, 163–174. 10.1016/j.biocon.2016.10.034

Navarro-Castilla, Á., & Barja, I. (2019). Stressful living in lower-quality habitats? Body mass, feeding behavior and physiological stress levels in wild wood mouse populations. Integrative Zoology, 14(1), 114–126. 10.1111/1749-4877.12351

Norman, A. F., Regnery, R., Jameson, P., Greene, C., & Krause, D. C. (1995). Differentiation of Bartonella-like isolates at the species level by PCR-restriction fragment length polymorphism in the citrate synthase gene. Journal of Clinical Microbiology, 33(7), 1797–1803. 10.1128/jcm.33.7.1797-1803.1995

Parker, K. L., Barboza, P. S., & Gillingham, M. P. (2009). Nutrition integrates environmental responses of ungulates. Functional Ecology, 23(1), 57–69. 10.1111/j.1365-2435.2009.01528.x

Perez, G., Bastian, S., Chastagner, A., Agoulon, A., Rantier, Y., Vourc’h, G., Plantard, O., & Butet, A. (2020). Relationships between landscape structure and the prevalence of two tick-borne infectious agents, Anaplasma phagocytophilum and Borrelia burgdorferi sensu lato, in small mammal communities. Landscape Ecology, 35(2), 435–451. 10.1007/s10980-019-00957-x

Plat, N., Allan, E., Jouveau, S., Bonnard, O., Rivoal, J.-B., Dudit, J., Barlier, T., Charbonnier, Y., Sannier, M., Garcia-Celada, I., van Halder, I., & Jactel, H. (2026). The role of broadleaved hedgerows for biodiversity conservation in pine plantation landscape. Biodiversity and Conservation, 35(2), 66. 10.1007/s10531-026-03273-4

Plat, N., Charbonnier, Y., Garcia-Celada, I., Sannier, M., Bonnard, O., Rivoal, J.-B., Ribot, T., Coupart, T., van Halder, I., & Jactel, H. (2025). Functional role of hedgerows and landscape composition in controlling a forest insect pest. Landscape Ecology, 40(8), 152. 10.1007/s10980-025-02170-5

Plowright, R. K., Reaser, J. K., Locke, H., Woodley, S. J., Patz, J. A., Becker, D. J., Oppler, G., Hudson, P. J., & Tabor, G. M. (2021). Land use-induced spillover: A call to action to safeguard environmental, animal, and human health. The Lancet Planetary Health, 5(4), e237–e245. 10.1016/S2542-5196(21)00031-0

Portillo, A., Santibáñez, P., Palomar, A. M., Santibáñez, S., & Oteo, J. A. (2018). ‘*Candidatus* Neoehrlichia mikurensis’ in Europe. New Microbes and New Infections, 22, 30–36. 10.1016/j.nmni.2017.12.011

Pradel, J., Bouilloud, M., Loiseau, A., Piry, S., Galan, M., Artige, E., Castel, G., Ferrero, J., Gallet, R., Thuel, G., Vieira, N., & Charbonnel, N. (2022). Small terrestrial mammals (Rodentia and Soricomorpha) along a gradient of forest anthropisation (reserves, managed forests, urban parks) in France. Biodiversity Data Journal, 10, e95214. 10.3897/bdj.10.e95214

Prist, P. R., Siliansky de Andreazzi, C., Vidal, M. M., Zambrana-Torrelio, C., Daszak, P., Carvalho, R. L., & Tambosi, L. R. (2023). Promoting landscapes with a low zoonotic disease risk through forest restoration: The need for comprehensive guidelines. Journal of Applied Ecology, 60(8), 1510–1521. 10.1111/1365-2664.14442

Randolph, S. E. (2004). Tick ecology: Processes and patterns behind the epidemiological risk posed by ixodid ticks as vectors. Parasitology, 129(S1), S37–S65. 10.1017/S0031182004004925

Renesto, P., Gouvernet, J., Drancourt, M., Roux, V., & Raoult, D. (2001). Use of rpoB Gene Analysis for Detection and Identification of Bartonella Species. Journal of Clinical Microbiology, 39(2), 430–437. 10.1128/JCM.39.2.430-437.2001

Riquelme, M., Salgado, R., Simonetti, J. A., Landaeta-Aqueveque, C., Fredes, F., & Rubio, A. V. (2021). Intestinal Helminths in Wild Rodents from Native Forest and Exotic Pine Plantations (Pinus radiata) in Central Chile. Animals, 11(2), 384. 10.3390/ani11020384

Rubio, A. V., Fredes, F., & Simonetti, J. A. (2019). Exotic Pinus radiata Plantations do not Increase Andes Hantavirus Prevalence in Rodents. EcoHealth, 16(4), 659–670. 10.1007/s10393-019-01443-1

Sánchez, C. A., Becker, D. J., Teitelbaum, C. S., Barriga, P., Brown, L. M., Majewska, A. A., Hall, R. J., & Altizer, S. (2018). On the relationship between body condition and parasite infection in wildlife: A review and meta-analysis. Ecology Letters, 21(12), 1869–1884. 10.1111/ele.13160

Sanderson, C. E., & Alexander, K. A. (2020). Unchartered waters: Climate change likely to intensify infectious disease outbreaks causing mass mortality events in marine mammals. Global Change Biology, 26(8), 4284–4301. 10.1111/gcb.15163

Santé publique France. (2026). Épidémiologie de la leptospirose en France en 2025. Données de la déclaration obligatoire: Édition nationale. Bulletin (p. 11) [Rapport]. Santé publique France.

Shah, H. A., Huxley, P., Elmes, J., & Murray, K. A. (2019). Agricultural land-uses consistently exacerbate infectious disease risks in Southeast Asia. Nature Communications, 10, 4299. 10.1038/s41467-019-12333-z

Singh, C., & Persson, U. M. (2026). Global patterns of commodity-driven deforestation and associated carbon emissions. Nature Food, 7(2), 138–151. 10.1038/s43016-026-01305-4

Smith, K. F., Goldberg, M., Rosenthal, S., Carlson, L., Chen, J., Chen, C., & Ramachandran, S. (2014). Global rise in human infectious disease outbreaks. Journal of The Royal Society Interface, 11(101), 20140950. 10.1098/rsif.2014.0950

Speakman, J. R. (2007). The physiological costs of reproduction in small mammals. Philosophical Transactions of the Royal Society B: Biological Sciences, 363(1490), 375–398. 10.1098/rstb.2007.2145

Spitz, F., Le Louarn, H., Poulet, A., & Dassonville, B. (1974). Standardisation des piégeages en ligne pour quelques espèces de rongeurs. Revue d’Écologie (La Terre et La Vie), 28(4), 564–578. 10.3406/revec.1974.4854

Stoddard, R. A., Gee, J. E., Wilkins, P. P., McCaustland, K., & Hoffmaster, A. R. (2009). Detection of pathogenic *Leptospira* spp. Through TaqMan polymerase chain reaction targeting the LipL32 gene. Diagnostic Microbiology and Infectious Disease, 64(3), 247–255. 10.1016/j.diagmicrobio.2009.03.014

Sweeny, A. R., Clerc, M., Pontifes, P. A., Venkatesan, S., Babayan, S. A., & Pedersen, A. B. (2021). Supplemented nutrition decreases helminth burden and increases drug efficacy in a natural host–helminth system. Proceedings of the Royal Society B: Biological Sciences, 288(1943), 20202722. 10.1098/rspb.2020.2722

Tattersall, F. H., Macdonald, D. W., Hart, B. J., Manley, W. J., & Feber, R. E. (2001). Habitat use by wood mice (Apodemus sylvaticus) in a changeable arable landscape. Journal of Zoology, 255(4), 487–494. 10.1017/S095283690100156X

Telfer, S., Begon, M., Bennett, M., Bown, K. J., Burthe, S., Lambin, X., Telford, G., & Birtles, R. (2007). Contrasting dynamics of Bartonella spp. in cyclic field vole populations: The impact of vector and host dynamics. Parasitology, 134(3), 413–425. 10.1017/S0031182006001624

Telfer, S., Clough, H. E., Birtles, R. J., Bennett, M., Carslake, D., Helyar, S., & Begon, M. (2007). Ecological Differences and Coexistence in a Guild of Microparasites: Bartonella in Wild Rodents. Ecology, 88(7), 1841–1849. 10.1890/06-1004.1

Tikhonov, G., Opedal, Ø. H., Abrego, N., Lehikoinen, A., de Jonge, M. M. J., Oksanen, J., & Ovaskainen, O. (2020). Joint species distribution modelling with the r-package Hmsc. Methods in Ecology and Evolution, 11(3), 442–447. 10.1111/2041-210X.13345

Unnsteinsdottir, E. R., & Hersteinsson, P. (2011). Effects of contrasting habitats on population parameters and diet of Apodemus sylvaticus (Rodentia) in south-estern Iceland. 75(1), 13–21. 10.1515/mamm.2010.068

Vanneste, T., Govaert, S., Spicher, F., Brunet, J., Cousins, S. A. O., Decocq, G., Diekmann, M., Graae, B. J., Hedwall, P.-O., Kapás, R. E., Lenoir, J., Liira, J., Lindmo, S., Litza, K., Naaf, T., Orczewska, A., Plue, J., Wulf, M., Verheyen, K., & De Frenne, P. (2020). Contrasting microclimates among hedgerows and woodlands across temperate Europe. Agricultural and Forest Meteorology, 281, 107818. 10.1016/j.agrformet.2019.107818

Vanroy, T., Baeten, L., Martel, A., Catfolis, B., Fonville, M., Lens, L., Pasmans, F., Sprong, H., Strubbe, D., Verbrugghe, E., & Verheyen, K. (2025). Influence of forest structural complexity on small mammal body condition and its impact on tick burden and pathogen prevalence. Parasites & Vectors, 18(1), 227. 10.1186/s13071-025-06874-0

Vié, J.-C., Hilton-Taylor, C., & Stuart, S. N. (Eds). (2009). Wildlife in a changing orld. IUCN. 10.2305/IUCN.CH.2009.17.en

Voutilainen, L., Savola, S., Kallio, E. R., Laakkonen, J., Vaheri, A., Vapalahti, O., & Henttonen, H. (2012). Environmental Change and Disease Dynamics: Effects of Intensive Forest Management on Puumala Hantavirus Infection in Boreal Bank Vole Populations. PLOS ONE, 7(6), e39452. 10.1371/journal.pone.0039452

Warton, D. I., Blanchet, F. G., O’Hara, R. B., Ovaskainen, O., Taskinen, S., Walker, S. C., & Hui, F. K. C. (2015). So Many Variables: Joint Modeling in Community Ecology. Trends in Ecology & Evolution, 30(12), 766–779. 10.1016/j.tree.2015.09.007

Wolton, R. J. (1985). The ranging and nesting behaviour of Wood mice, Apodemus sylvaticus (Rodentia: Muridae), as revealed by radio-tracking. Journal of Zoology, 206(2), 203–222. 10.1111/j.1469-7998.1985.tb05645.x

Zarea, A. A. K., Laidoudi, Y., Odigie, A. E., Mrenoshki, D., Iulietto, M. F., Greco, G., & Condoleo, R. (2025). Estimated prevalence and genetic diversity of *Bartonella* species in small mammals in Europe: Systematic review and meta-analysis. One Health, 21, 101263. 10.1016/j.onehlt.2025.101263

Zawada, S. G., Fricken, M. E. von, Weppelmann, T. A., Sikaroodi, M., & Gillevet, P. M. (2020). Optimization of tissue sampling for Borrelia burgdorferi in white-footed mice (Peromyscus leucopus). PLOS ONE, 15(1), e0226798. 10.1371/journal.pone.0226798

Zintl, A., McManus, A., Galan, M., Diquattro, M., Giuffredi, L., Charbonnel, N., Gray, J., Holland, C., & Stuart, P. (2023). Presence and identity of *Babesia microti* in Ireland. Ticks and Tick-Borne Diseases, 14(6), 102221. 10.1016/j.ttbdis.2023.102221

Zuk, M., & McKean, K. A. (1996). Sex differences in parasite infections: Patterns and processes. International Journal for Parasitology, 26(10), 1009–1023. 10.1016/S0020-7519(96)80001-4

