## Supplementary Methods for "Intensive pine plantation management shapes infectious agent communities in the generalist rodent *Apodemus sylvaticus*, with consequences for zoonotic hazard"

### Species characterization of Bartonella spp. using *gltA* and *rpoB* metabarcoding

We have prepared Illumina libraries using 2-step PCR following the protocol described in Galan et al 2018. We performed two technical replicates on each DNA extract of spleen samples in order to control PCR stochasticity for rare DNA targets and to validate the positive results (Robasky et al 2014). Negative controls for extraction (NCext, 1 per DNA extraction session), PCR (1 NCpcr) and indexing (1 NCindex) were included on each 96-PCR microplates.

#### a) Target specific amplification (PCR1) of *gltA* gene:

A first amplification step of 338 bp *gltA* fragment was carried out using primers (BhCS.781p 5’-GGGGACCAGCTCATGGTGG-3’ and BhCS.1137n	5’-AATGCAAAAAGAACAGTAAACA-3’) described in Norman et al 1995 and modified by 5' end overhang Illumina linker. The PCR reactions were carried out in a final volume of 10 µL containing 5 µL of Multiplex Master Mix (Qiagen, Hilden, Germany), 0.5 μM of each primer and 1.5 µL of DNA. The PCR conditions for this first step consisted of initial denaturation at 95°C for 15 min followed by 40 cycles of denaturation at 94°C for 30 s, hybridization at 50°C for 45 s and extension at 72°C for 2 min, and a final extension step at 72°C for 10 min.

***gltA* primers :**

BhCS.781p-Illumina	TCGTCGGCAGCGTCAGATGTGTATAAGAGACAGGGGGACCAGCTCATGGTGG

BhCS.1137n-Illumina	GTCTCGTGGGCTCGGAGATGTGTATAAGAGACAGAATGCAAAAAGAACAGTAAACA

#### Target specific amplification (PCR1) of the *rpoB* gene:

A first amplification step of 333 bp *rpoB* fragment was carried out using primers (BartRpoB.1673-F 5’-CGCATTATGGTCGTATTTGTCC-3’ and BartRpoB.2006-R	5’-GCACGATTCGCATCATCATTTTCC-3’) described in Renesto et al 2001 (but with reverse complemented sequences) and modified by 5' end overhang Illumina linker. The PCR reactions were carried out in a final volume of 10 µL containing 5 µL of Multiplex Master Mix (Qiagen, Hilden, Germany), 0.5 μM of each primer and 1.5 µL of DNA. The PCR conditions for this first step consisted of initial denaturation at 95°C for 15 min followed by 40 cycles of denaturation at 94°C for 30 s, hybridization at 50°C for 45 s and extension at 72°C for 2 min, and a final extension step at 72°C for 10 min.

***rpoB* primers :**

BartRpoB.1673-F-Illumina	TCGTCGGCAGCGTCAGATGTGTATAAGAGACACGCATTATGGTCGTATTTGTCC

BartRpoB.2006-R-Illumina	GTCTCGTGGGCTCGGAGATGTGTATAAGAGACAGCACGATTCGCATCATCATTTTCC

#### c) Dual Indexing (PCR2):

A second PCR step was performed to add individual-specific multiplexing tags (called index i5 and index i7), consisting of short 8 bp sample-specific sequences (Kozich et al 2013) and the Illumina adapters (called P5 and P7) at the 5' ends of each amplified DNA fragment to the first PCR1. As all the PCR2 products were mixed together (multiplexing) for MiSeq sequencing, dual-indexes made it possible to identify the origin of the sequences and reassign them to each sample (demultiplexing). Each pair of i5 and i7 indexes were used for a unique PCR replicate (i.e. unique dual indexes) to enable that libraries were sequence and demultiplex. This second PCR was carried out in a total volume of 10 µL containing 5 µL of Multiplex kit (Qiagen, Germany), 0.7 μM of each i5 and i7 primers, and 2 µL of products from the first PCR for each sample. PCR conditions consisted of an initial denaturation step at 95°C for 15 min followed by 8 cycles of denaturation at 95°C for 40 s, hybridization at 55°C for 45 s and extension at 72°C for 2 min, and a final extension step at 72°C for 10 min.

Forward indexed i5 primer PCR2 sequence (i5 & i7 sequences are listed in Kozich et al 2013):

AATGATACGGCGACCACCGAGATCTACAC<i5>TCGTCGGCAGCGTC

Reverse indexed i7 primer PCR2 sequence:

CAAGCAGAAGACGGCATACGAGAT<i7>GTCTCGTGGGCTCGG

#### d) Pooling, size selection & sequencing:

The volume-to-volume mixes (one for each gene) of all the PCR products (4 µL per sample) were screened based on fragment length resulting from excision on 1.25% agarose gel. The pools of specific PCR products were cut to the expected sizes (516 bp for both *gltA* and *rpoB* PCR products, corresponding to the size of the amplicon, including the gene-specific primers, sequencing primers, indexes and adaptors) under UV light. The resulting libraries were purified with the PCR purification kit (Macherey-Nagel, Germany), quantified by quantitative PCR (KAPA kit, Kapa Biosciences), then sequenced on a MiSeq sequencer with a 500v2 kit (Illumina, USA).

### References:

Galan, M., Pons, J.-B., Tournayre, O., Pierre, É., Leuchtmann, M., Pontier, D., & Charbonnel, N. (2018). Metabarcoding for the parallel identification of several hundred predators and their prey: Application to bat species diet analysis. Molecular Ecology Resources, 18, 474–489. <https://doi.org/10.1111/1755-0998.12749>

Kozich, J. J., Westcott, S. L., Baxter, N. T., Highlander, S. K., & Schloss, P.D. (2013). Development of a dual-index sequencing strategy and curation pipeline for analyzing amplicon sequence data on the MiSeq Illumina sequencing platform. Applied and Environmental Microbiology, 79(17), 5112–5120. <https://doi.org/10.1128/aem.01043-13>

Robasky, K., Lewis, N. E., & Church, G. M. (2014). The role of replicates for error mitigation in next-generation sequencing. Nature Reviews Genetics, 15(1), 56–62. <https://doi.org/10.1038/nrg3655>
