## Supplementary Figure for "Intensive pine plantation management shapes infectious agent communities in the generalist rodent *Apodemus sylvaticus*, with consequences for zoonotic hazard"

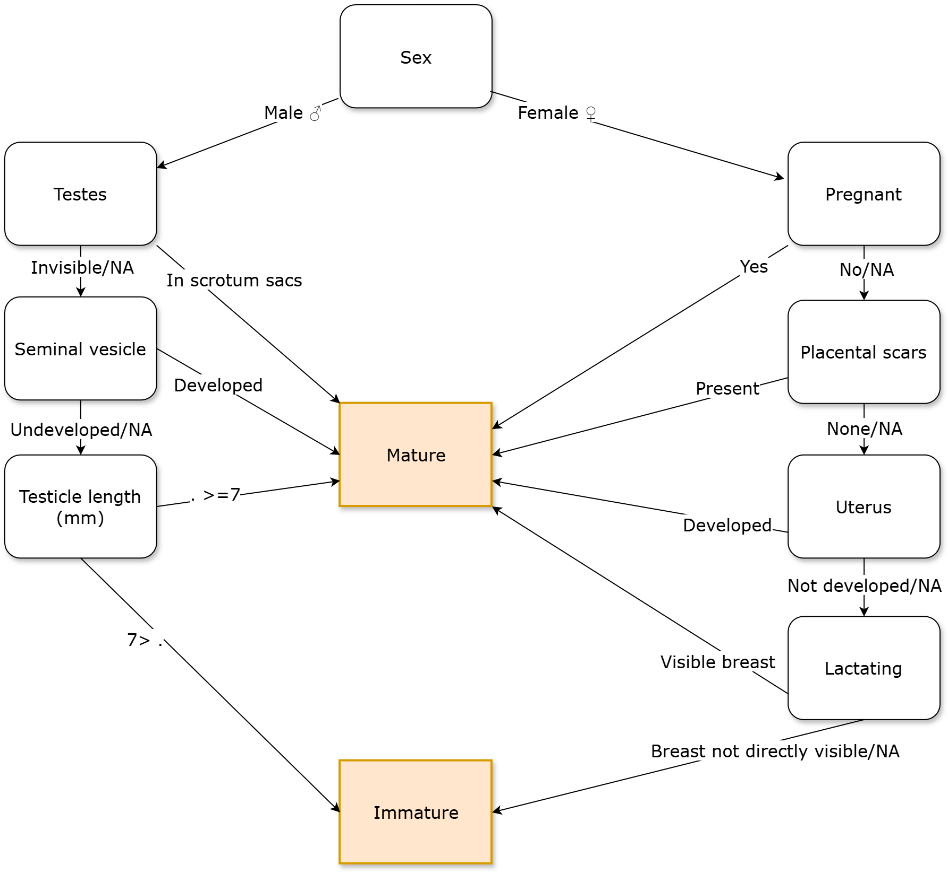

**FIGURE S1. Decision tree for determining the sexual maturity of *Apodemus sylvaticus*. Based on external and internal reproductive characteristics.**

For males, a testis length threshold of 7 mm was empirically determined from our dataset as the value that best separated individuals of confirmed sexual maturity from the remaining males, thereby resolving ambiguous cases.

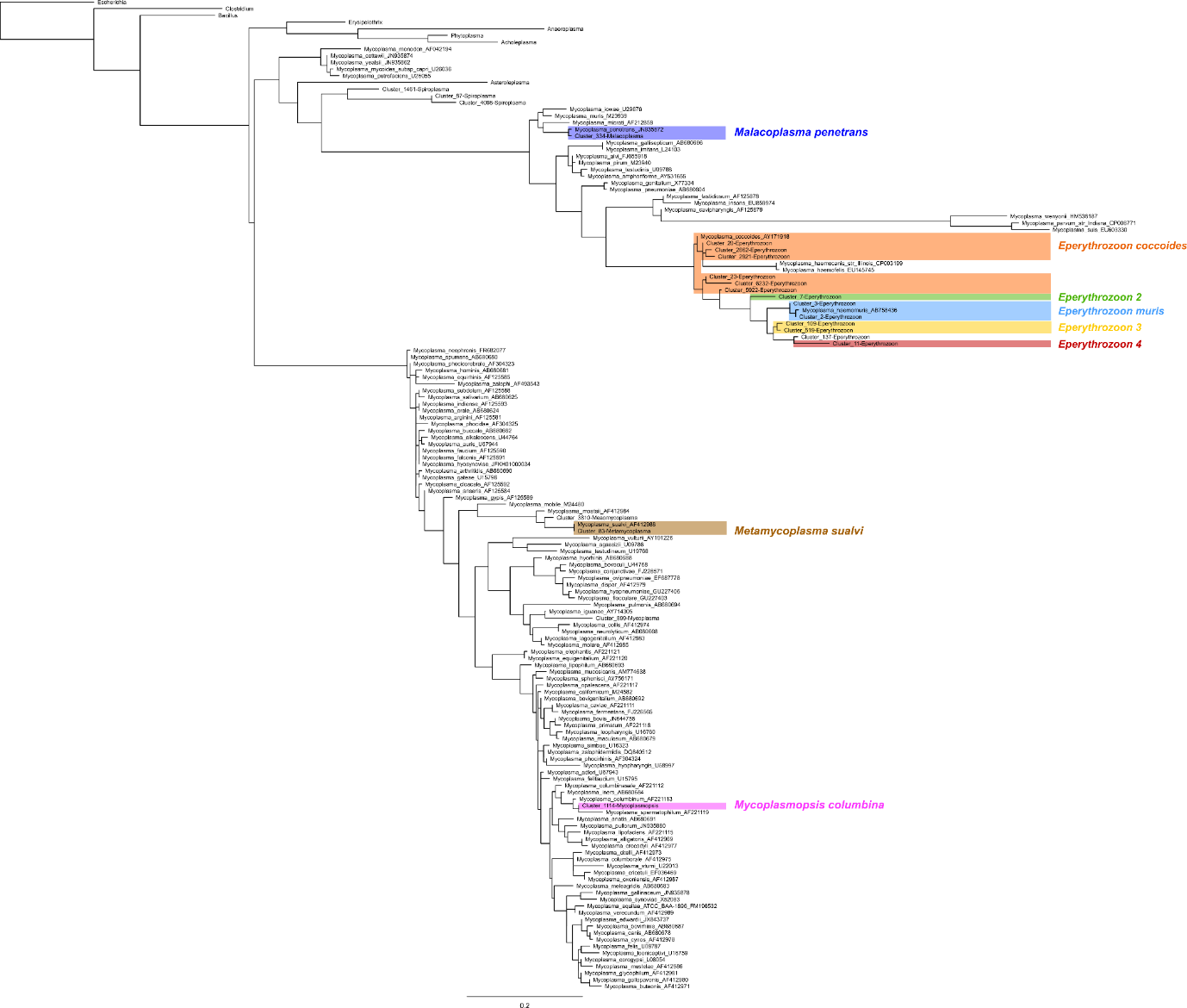

**FIGURE S2. Maximum likelihood phylogenetic tree of *Mycoplasmatota* based on 16S rRNA V4 sequences.**

FASTA sequences were clustered into operational taxonomic units (OTUs) using Swarm (*d* = 1) within the FROGS pipeline (Escudié et al., 2018). OTUs were initially assigned a taxonomy using the SILVA database (release 138.2, Pintail100). OTUs affiliated to the *Mycoplasmataceae* family were selected for further analyses and are labeled as “Cluster_XX” in the figure. Representative OTU sequences were aligned with reference sequences using MAFFT, and the maximum-likelihood phylogeny was inferred with IQ-TREE using the GTR+I+G nucleotide substitution model. OTU identities were further validated by BLAST searches against the NCBI Nucleotide database, allowing confirmation of taxonomic assignments or grouping of closely related sequences for downstream analyses. Initial SILVA taxonomy was subsequently updated according to the revised taxonomy proposed by Gupta et al. (2018). Colored clusters indicate the main phylogenetic groups: pink, sequences closely related to *Mycoplasmopsis columbina*; brown, *Metamycoplasma sualvi*; dark blue, *Metamycoplasma penetrans*; orange, *Eperythrozoon coccoides*; light blue, *Eperythrozoon muris*. Yellow, green, and red clusters correspond to distinct haemoplasma lineages. Although these sequences grouped with other *Eperythrozoon* spp., they were too divergent to allow confident species-level assignment and were therefore treated as three separate *Eperythrozoon* groups.

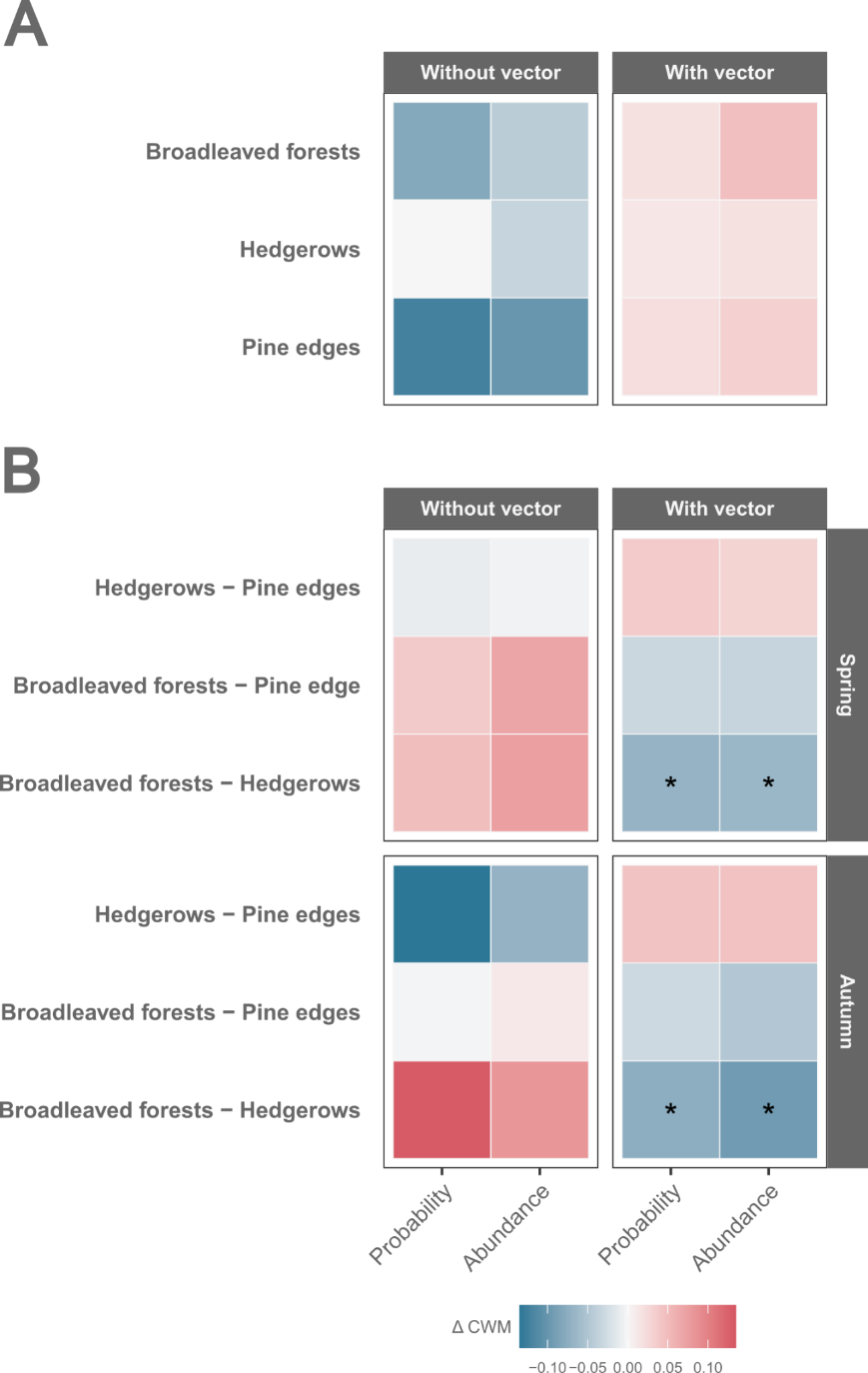

**FIGURE S3. Seasonal and habitat effects on predicted community-weighted mean (CWM) transmission traits inferred from HMSC models.**

**A.** Pairwise differences in predicted marginal CMW between spring and autumn (spring - autumn) for each habitat type (rows), transmission trait (panels), and HMSC modelling approach (columns; Probability or Abundance). **B.** Pairwise differences in predicted marginal CMW for each habitat comparison (rows), season (row panels), transmission trait (column panels), and HMSC approach (columns; Probability or Abundance). Pairwise differences were calculated for each posterior sample as the difference between predicted marginal CMW. Asterisks indicate supported contrasts (posterior support, PS ≥ 0.95), where PS is the proportion of posterior samples sharing the same sign as the posterior mean. Predicted CWM differences are expressed on the same scale for both modelling approaches.

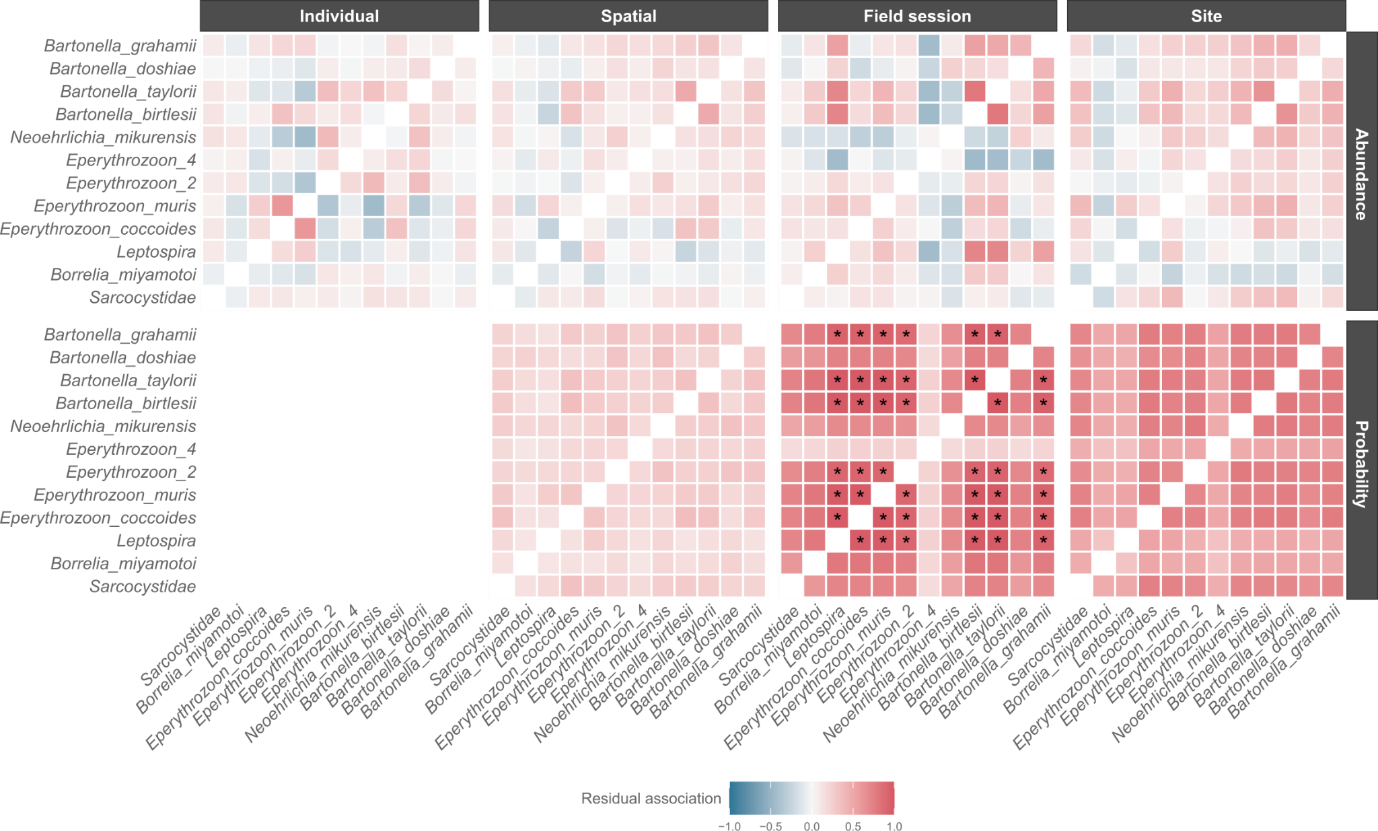

**FIGURE S4. Residual associations variance-covariance matrix between infectious taxa**

Residual association (Ω) between taxa for the infection probability and infected-host abundance approaches (row panels) and every random effect included (column panels). Asterisks denote supported contrasts (posterior support, PS ≥ 0.95), where PS is the proportion of posterior samples sharing the same sign as the posterior mean.
