## Supplementary Tables captions for "Intensive pine plantation management shapes infectious agent communities in the generalist rodent *Apodemus sylvaticus*, with consequences for zoonotic hazard"

**TABLE S1. Biologically relevant hypotheses tested to explain variation in infectiosome responses in relation to host characteristics and environmental determinants.**

**A.** Candidate hypotheses evaluated using generalized linear mixed models (GLMMs) for community-level response measured as infectious taxon richness within individual hosts. **B.** Candidate hypotheses evaluated using hierarchical modelling of species communities (HMSC) for taxon-specific response measured as infection probability of each taxon. **C.** Candidate hypotheses evaluated using HMSC for taxon-specific response measured as the abundance of infected individuals for each taxon. The same hypothesis-driven candidate model framework was applied across the GLMM and HMSC analyses to ensure consistency in model selection and biological interpretation.

**TABLE S2. Number of distinct *Apodemus sylvaticus* individuals at each site-session**

Number of individuals captured, and corresponding number of individuals from which tissues were collected for infectious agent screening.

**TABLE S3. Model selection for individual-level infectious taxon richness.**

Candidate models were compared using the corrected Akaike information criterion (AICc).

**TABLE S4. Performance and convergence metrics of candidate HMSC models**

**A.** Performance and convergence metrics for models fitted to estimate the probability of infection. **B.** Performance and convergence metrics for models fitted to estimate infected-host abundance. Reported values correspond to averages across all infectious taxa included in the analyses (n = 12).

**TABLE S5. Performance metrics of each candidate HMSC models for each infectious taxon**

**A.** Performance metrics for each taxon and models fitted to estimate the probability of infection. **B.** Performance metrics for each taxon and models fitted to estimate infected-host abundance.
